# Neural crest cells give rise to non-myogenic mesenchymal tissue in the adult murid ear pinna

**DOI:** 10.1101/2023.08.06.552195

**Authors:** Robyn S. Allen, Shishir K. Biswas, Ashley W. Seifert

## Abstract

Despite being a major target of reconstructive surgery, development of the external ear pinna remains poorly studied. As a craniofacial organ highly accessible to manipulation and highly conserved among mammals, the ear pinna represents a valuable model for the study of appendage development and wound healing in the craniofacial complex. Here we provide a cellular characterization of late gestational and postnatal ear pinna development in *Mus musculus* and *Acomys cahirinus* and demonstrate that ear pinna development is largely conserved between these species. Using *Wnt1-cre;ROSA^mT/mG^* mice we find that connective tissue fibroblasts, elastic cartilage, dermal papilla cells, dermal sheath cells, vasculature, and adipocytes in the adult pinna are derived from cranial crest. In contrast, we find that skeletal muscle and hair follicles are not derived from neural crest cells. Cellular analysis using the naturally occurring *short ear* mouse mutant shows that elastic cartilage does not develop properly in distal pinna due to impaired chondroprogenitor proliferation. Interestingly, while chondroprogenitors develop in a mostly continuous sheet, the boundaries of cartilage loss in the *short ear* mutant strongly correlate with locations of vasculature-conveying foramen. Concomitant with loss of elastic cartilage we report increased numbers of adipocytes, but this seems to be a state acquired in adulthood rather than a developmental abnormality. In addition, chondrogenesis remains impaired in the adult mid-distal ear pinna of these mutants. Together these data establish a developmental basis for the study of the ear pinna with intriguing insights into the development of elastic cartilage.

## INTRODUCTION

Cranio-facial tissues are a major target of reconstructive surgeries and tissue bioengineering for the repair of developmental defects and injuries (Cubitt et al., 2019; Habal, 2004; Jia et al., 2022; Schantz et al., 2012). As a highly accessible and anatomically well conserved cranio-facial tissue (Chiu et al., 2017), the external ear pinna represents a promising model for the study of both development and wound healing. Not surprisingly, however, little is known about the molecular regulation of external ear pinna development and the origin of the cells that give rise to the pinna proper. While detailed development of the inner and middle ear has been well-studied in mammals (Anthwal and Thompson, 2016; Whitfield, 2015), external ear development is poorly understood.

Available developmental data of the ear pinna comes from limited observations of human and mouse embryos (Cox et al., 2014; Green and Green, 1942; Hashimoto et al., 2021; Honkura et al., 2020; Hunter and Yotsuyanagi, 2005; Mallo and Gridley, 1996; Minoux et al., 2013; Theiler and Sweet, 1986b). In approximately the fourth week of human development, six nodular tissue swellings known as the Hillocks of His appear on the first and second pharyngeal arches and these eventually remodel to form the auricle or pinna ((rev. in Hunter and Yotsuyanagi, 2005)). While similar patterns have been observed in early histological studies of mutant mice, development of the structures that make up the pinna (external auditory meatus, scapha, helix, anti-helix, tragus and anti-tragus) are altered by specific genetic mutations (Green and Green, 1942; Theiler and Sweet, 1986a). More recent molecular studies in mice have revealed a complex genetic contribution to ear pinna development (Cox et al., 2014; Palmer et al., 2016) and inactivation of *Hoxa2* leads to a complete absence of the ear pinna (Mallo and Gridley, 1996; Rijli et al., 1993). The most comprehensive study to date demonstrated that the ear pinna is derived from *Hoxa2*-positive neural crest cells from the second pharyngeal arch only, rather than from the first and the second arches (Minoux et al., 2013). Furthermore, *Hoxa2* controls *Bmp4* and *Bmp5* expression by binding to upstream non-coding regions and Bmp-signaling is known to play a role during pinna development (Minoux et al., 2013). However, these studies only examined the ear pinna *in utero* (up to E18.5).

In addition to developmental studies, the ear pinna also represents a powerful model for studying musculoskeletal wound healing and regeneration. Wounding studies in cranial neural crest derived dermis shows that it has unique gene expression and extracellular matrix deposition along with increased migratory capacity compared to trunk skin (Usansky et al., 2021). These features may contribute to scar resistance in the cranio-facial complex. In addition, the external ear pinna is a synapomorphy among Therian mammals and as a structure it exhibits wide phenotypic variation (Manley, 2012). Despite variation in shape and size, the general tissue architecture of the pinna appears highly conserved with hair, skin, connective tissue and skeletal muscle surrounding an elastic cartilage frame (Ekdale, 2016; Gawriluk et al., 2016). This conserved anatomy, and the accessibility of the external ear pinna, has made it possible to screen many mammalian species for regenerative ability using a simple ear punch assay (Gawriluk et al., 2016; Williams-Boyce and Daniel, 1986). This led to the discovery of a highly regenerative rodent, the spiny mouse (*Acomys*) which are capable of regenerating full-thickness skin injuries and the complex musculoskeletal architecture of the external ear pinna in response to complete excision with a punch biopsy (Gawriluk et al., 2016; Matias Santos et al., 2016; Seifert et al., 2012). Further study of ear pinna development would provide insight into the regenerative potential of these animals and cellular mechanisms that may be curtailed in non-regenerating rodents.

To extend previous work and provide a foundation to better understand normal ear pinna development, developmental malformations, and pinna regeneration, we sought to characterize and compare post-natal development of the ear pinna in *Acomys cahirinus* and *Mus musculus*. Using this comparison, we show that tissue development in the ear pinna follows the same general pattern in these two species. Using genetic tools available for *Mus*, we show that neural crest cells contribute to most adult pinna tissues except for (1) the epidermis and epidermally-derived structures, (2) skeletal muscle and (3) a sparse population of cells found throughout the connective tissue. Finally, we used the naturally occurring *short ear* mutation to show that Bmp-signaling is required for the proliferation of chondroprogenitors in the mid-distal ear. Further, structural changes in the ear pinna resulting from defective cartilage formation appear to promote adipogenesis. Interestingly, chondrogenesis in the mid-distal ear continues to be defective even in adulthood in the absence of Bmp5.

## METHODS

### Animals and tissue collection

Outbred *Mus musculus* (Swiss Webster, Charles River and ND4, Envigo), *Wnt1-Cre* (Jax Strain 022137), *ROSA^mT/mG^*(JAX strain 007676), and *SEA/Gn* (JAX Strain 000644) animals were maintained in static microisolator cages and *Acomys cahirinus* were maintained as previously described(Haughton et al., 2016). *Mus musculus* breeding pairs (1 male and 1 female of breeding age) were checked for plugs every morning before 9am. Ear pinnae were harvested from mice at E20.5, P0, P1 and every other day afterwards until weaning at P21. Sample sizes for time points shown in Figure 2 are: E20.5 n=5, P1 n=5, P5 n=5, P9 n=5, P15 n=3, P21 n=3. *Acomys cahirinus* breeding cages were set up with 1 male to 3 females in large wire mesh cages. Since spiny mice do not plug, developmental stage of the embryonic time points was estimated based on embryonic characters including external genitalia, skin, limbs, etc. Tissue samples were collected from embryos estimated to be E20, E25, E30 and E35 and from post-natal time points at P0, P1, P5 and every five days after until P25. Sample sizes for time points shown in Figure 3 are: E20 n=1, E25 n=1. E35 n=1, P1 n=2, P5 n=2, P15 n=2. To trace neural crest cells *Wnt1-Cre* males were crossed to female *ROSA^mT/mG^*reporter mice. *Wnt1;ROSA^mT/mG^*pups were fostered to Swiss Webster mothers to avoid cannibalization. Tissue was collected at P11 (n=3) when all adult cellular compartments in the pinna had formed as well as at P50 (n=3) after the pinna had completely formed. Samples were set in 30% sucrose for 1 hour before embedding in OCT (TissueTek, Cat.# 4583) for cryosectioning without fixation. The *SEA/Gn* strain segregates the *short ear* allele, a single base point mutation in the *bone morphogenetic 5* (*Bmp5*) protein. The result of this mutation is a premature stop codon where the truncated protein lacks the 3’ carboxy signaling motif. *Short ear* mice were maintained by crossing *short ear^-/+^* to *short ear^-/-^*, resulting in Mendelian litters of heterozygous and null animals. Both breeding strategies (null female x het male; het female x null male) were used with no apparent difference in breeding efficiency. Tissue was collected from null and heterozygous offspring (n=3 each for paraffin and cryosectioning) at weening and were fixed overnight. All animal procedures were approved by the University of Kentucky Institutional Animal Care and Use Committee (IACUC) under protocol 2013-1119.

### Genotyping

*Shot ear Mus* were genotyped phenotypically at weaning. *Bmp5*^se/se^ (referred to as Bmp5^-/-^) mutant mice have characteristic truncated and malformed ears, while *Bmp*5^+/se^ (referred to as Bmp5^+/-^) develop normally. Prenatal Bmp5^+/-^ and ^-/-^ mice were genotyped using sanger sequencing according to protocol recommendations by Jackson Laboratories. DNA was isolated from tail tips using overnight digestion at 55°C in lysis buffer (100mM Tris-Cl, 5 mM EDTA, 0.2% SDS, and 200µg/mL Proteinase K), followed by 10-minute heat inactivation at 90°C. DNA was PCR amplified to enrich for a *short ear* mutation containing segment of the *Bmp5* gene using the following primers (F – GAACCATTTCACCAGCTCCT, R – GGAGGCATTACAAAGAGTTTCG). PCR products were purified using gel electrophoresis and gel purification using a QiAquick gel extraction kit (Qiagen, Cat. #28704). Purified PCR products were sent for sanger sequencing using the same primers as used in the RCR amplification step. *Bmp5*^-/-^ mutants were identified based on the presence of the *short ear* C to G substitution.

### Whole mount Alcian blue staining

Whole ears were harvested from adult individuals and fixed overnight in 10% neutral buffered formalin. Fixed tissues were rinsed three times in phosphate buffered saline for 10 minutes each and then incubated in 2.5% trypsin solution at 37°C for 30minutes. Tissues were then washed for 10 minutes in phosphate buffered saline and the dorsal tissue (epidermis, dermis and muscle) layer was peeled away from the underlying cartilage. Partially intact ears were dehydrated by rinsing three times in 70% ethanol for 10 minutes each and then transferred to 100% ethanol for overnight incubation at 4°C. Dehydrated ears were then incubated with 0.2% Alcian blue solution for 8 hours at 25°C, followed by clearing solution (1% KOH/10% glycerol in H_2_O) overnight at 25°C. Tissues were then transferred to storage solution (50% glycerol/50% ethanol) for imaging. Ventral epidermis and dermis were peeled away from the underlying cartilage leaving an intact sheet of cartilage tissue. Cartilage sheets were mounted in storage solution for imaging.

### Histology and Immunostaining

Harvested tissue was fixed overnight at 4°C in 10% neutral buffered formalin followed by three, ten-minute rinses in phosphate buffered saline and then 70% ethanol. Tissue was then embedded in paraffin blocks and sectioned at 5µm for histology and immunostaining. Masson’s Trichrome staining was performed using an American Mastertech Kit (Cat.# KTMTRPT) using paraffin sections. Alcian blue staining was performed using a solution of 1% w/v Alcian blue dissolved in 3% v/v acetic acid and counterstained with Brazilliant (Anantech, Cat.# NC9982671). Stained histological sections were mounted with XYL seal (Thermo Scientific, Cat.# 8312-4).

Immunohistochemistry was performed as previously described (Gawriluk et al., 2016). Primary antibodies used for immunostaining were anti-Tenascin-C (Tnc) (Millipore Cat.# AB19013), anti-Fatty acid binding protein 4 (Fabp4) (Abcam, Cat.# ab92501), anti-Myosin Heavy Chain 3 (Myh3) (Abcam, Cat.# ab124205), anti-Ionized calcium binding adapter molecule 1 (Iba1) (Wako, Cat.# 019-1971), anti-SRY Box Transcription Factor 9 (Sox9) (Abcam, Cat.# ab185230), and anti-Ki67 (Abcam, Cat.# ab15580). Secondary antibodies used were Alexafluor 594 Donkey-anti-Rabbit (Life Technologies, Cat.# A21207) and Cy5 AffiniPure Goat-anti-Rabbit (Jackson ImmunoResearch, Cat.# 711-606-152). Ki67 was additionally enhanced using biotin-streptavidin amplification with goat anti-rabbit biotinylated IgG (Vector Laboratories, Cat.# BA-1000) and Streptavidin conjugated Alexafluor 594 (Life Technologies, Cat.# 532356). Slides were mounted using ProLong Gold (Invitrogen, Cat. # P36934). Antigen retrieval was performed when immunostaining for Tenascin-C by exposure to Proteinase K for 2-5 minutes, for Iba1 by heat retrieval in Tris-EDTA ph9 buffer, and for Ki67 by heat retrieval in citrate buffer pH 6. Negative controls were run using an isotyped IgG specific to the primary species.

### Image collection and processing

Fluorescent and bright field images were collected using an Olympus BX53 fluorescent deconvolution microscope (Olympus America Inc). Bright field images were taken for whole mount, Masson’s trichrome and Alcian blue. Images of *Wnt1-cre;ROSA^mT/mG^*cryosectioned tissues were obtained using a Leica TCS SP Laser Scanning confocal microscope. Brightness and contrast were optimized using Adobe Photoshop as needed. Fluorescent images were taken in single channels in grayscale for immunostaining. Images were then pseudocolored and brightness, contrast and histogram maximum/minimum levels were optimized for each channel individually as needed and then overlaid.

### Histological Measurements and Statistics

Histological characteristics such as total ear thickness, cartilage thickness and number of hair follicles as well as Fabp4+ areas were measured using ImageJ software (National Institutes of Health, Bethesda, MD). Percentages for *Wnt1-* and Fabp4+ cells were calculated by hand-counting the total number of positive cells and dividing by the total number of Hoescht+ nuclei.

Student’s T-tests and One and Two-Way ANOVAs were performed as necessary on counts from histological images using JMP 12 (SAS Institute Inc., Cary, NC) or Prism (Graphpad Software, Boston, MA).

### Ear pinna Fibroblast isolation and culture

Ear pinna fibroblasts were isolated and cultured as previously described (Saxena et al., 2019). Briefly ear tissue was digested in 2.5% trypsin:dispase 1:1 solution for 30-60 minutes at 37°C. Ear tissue from *short ear* heterozygotes and null mutants were separated in to proximal 2/3 and distal 1/3 prior to digestion. Tissues were mechanically digested and then digested in collagenase. Following a final mechanical digestion, cells were washed and plated in complete DMEM with 10% FBS and 1% antibiotic-antimycotic solution. Cells were cultured on plastic tissue culture plates at 3% oxygen for 2 passages before undergoing chondrogenesis.

### In vitro chondrogenesis

In vitro chondrogenesis was performed according to previously established protocols (Culbert et al., 2014; De Ceuninck et al., 2004). Briefly, cells were harvested using 2.5% trypsin and washed. They were then suspended in a 2.5% alginate solution. The cell solution was passed through a 32-gauge needle into polymerization solution (102mM CaCl_2_, 10mM HEPES) resulting in the formation of 3D spheres approximately 20-30 mm in diameter. 3D spheres were washed in 0.9% NaCl and then chondrogenesis medium (High glucose DEMEM with 0.1µM Dexmethasone (Sigma, Cat.# D4902), 50µg/mL L-ascobate-2-phosphate, 40µg/mL L-proline, L-glutamine, 100µg/mL sodium pyruvate, 1:100 ITS+ culture supplement (Stemcell Technologies, Cat.# 07151), and 50ng/mL hrBMP4 (RandD Systems, Cat.# 314-BP)). Spheres were separated into individual wells on a 24 well plate and cultured in chondrogenesis medium for 21 days, with media changes every 2-3 days, before harvest. Cells were released from alginate spheres using dissolution solution (55mM EDTA, 10mM HEPES) for QPCR. Intact alginate spheres were fixed in 10% formalin and paraffin embedded for alcian blue staining.

### Quantitative PCR

Cells were collected in trizol and RNA was isolated using phenol chloroform purification and ethanol precipitation. RNA quality was assessed using gel electrophoresis. RNA concentration was measured using a nanodrop. cDNA was generated from purified RNA using the Sensifast cDNA synthesis kit (Cat. # BIO-65054). QPCR was performed using SyberGreen fastmix (Quanta, Cat. # 95072-012) in a Roche Lightcycler 96 system. The following primers were used for amplification of *Mus* cDNA: *sox9* (F – AGCCCTTTCAACCTGCAGC, R – TTCCTTTCCAGTCTTTGTGAGAACA), *collagen 2a* (*col2a*) (F – CACCCACATCACCCTCTGAC, R - GTAGGCTGGCTCCCATTCAG), and *aggrecan* (*acan*) (F – ACGAGGCAGACAGTACCTTG, R – CAGCCCTGGTTGGGATCAAT).

## RESULTS

### General tissue architecture of the ear pinna is well-conserved across murids

Murid rodents possess external ear pinnae with the same basic anatomy (Figure 1A). Using sexually mature (≥6 months) *Acomys cahirinus* and outbred laboratory *Mus musculus* (Swiss Webster and ND4 strains) as example murids, we compared general characteristics of the adult ear pinna (Figure 1B). First, we found the pinna was significantly thicker in *Acomys* versus *Mus* (413µm ± 52 versus 232µm ± 45; T=-5.23, *p*=0.002) as was the elastic cartilage (*Acomys*: 66µm ± 2 versus *Mus*: 47µm ± 6, T=-6.09, *p*=0.001). However, when we controlled for ear thickness, we found elastic cartilage in *Mus* occupied a larger proportion of the pinna compared to *Acomys* (*Acomys*: 0.162 ± 0.019 versus *Mus*: 0.202 ± 0.02 cartilage thickness/ear thickness, T=2.94, *p*=0.026). We also quantified hair follicle number and found that *Acomys* had significantly more hair follicles on the dorsal side of the pinna compared to *Mus* (6.75 follicles/mm versus 2.5 follicles/mm, T=-4.16, *p*=0.006) whereas there was no difference on the ventral portion of the pinna epidermis (1.2 follicles/mm versus 1 follicle/mm, p=0.62). Besides these gross anatomical differences, the overall cellular architecture of the adult ear pinna was nearly identical between species (Figure 1B). We further examined elastic cartilage in both species using whole mount alcian blue staining. Both species have a single sheet of elastic cartilage that spans the ear pinna (Figure 1C and D, Supplemental Figure 1). Interestingly, within this sheet, we observed foramen scattered throughout the elastic cartilage which were concentrated more distally. In *Mus*, the foramina appeared more densely in the posterior ear, with a prominent area of solid cartilage along the anterior edge (Figure 1C’), whereas in *Acomys* the foramina were more evenly distributed (Figure 1D’). Examination of ears from multiple individuals of each species suggests there is no regular pattern to these foramina.

**Figure 1.**
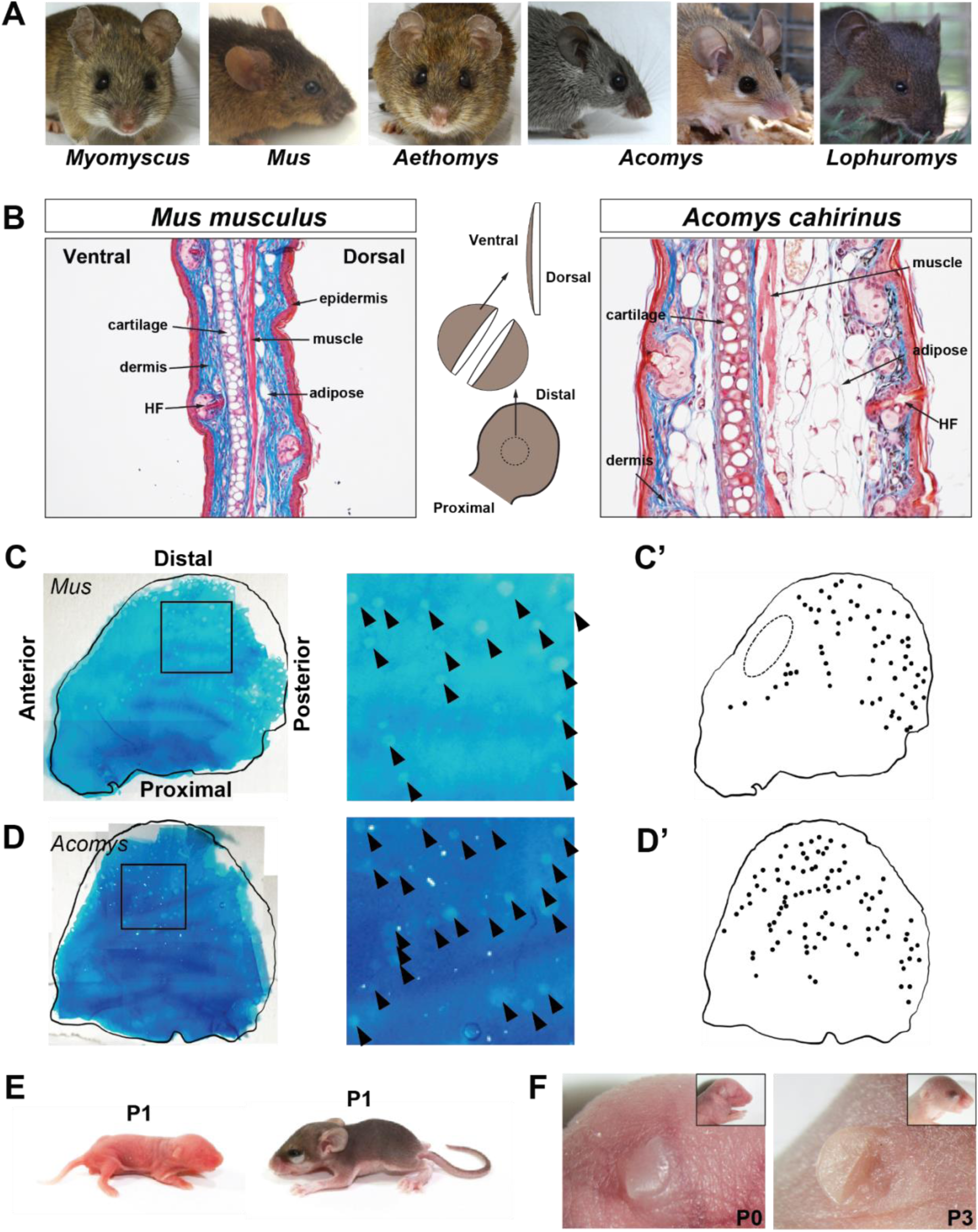
The structure of the external ear pinna is highly conserved among *murids*. (A) Examples of the ear pinna across multiple murid rodents. (B) Masson’s Trichrome tissue architecture of sexually mature (≥6 months) *Mus musculus* and *Acomys cahirinus* with schematic representation of proximodistal and dorsoventral orientation. In trichrome stained sections, nuclei are stained black, cytoplasm is stained pink/red and connective tissue and extracellular matrix are stained blue. (C-D) Whole mount alcian blue staining of adult *Mus* (C; n=2) and *Acomys* (D; n=1) ear pinna elastic cartilage. Arrowheads indicate individual foramen within the cartilage structure. (C’-D’) Schematic representation of foramina visible in the adult *Mus* (C’) and *Acomys* (D’) ear pinna elastic cartilage. Dotted oval indicates area of uninterrupted cartilage at the anterior edge. Ear outline and peripheral foramina locations were generated from partially intact ear pictures (see Supplemental Figure 1). (E) Newborn mouse (left) and spiny mouse (right). (F) Side view of a newborn (left; wildtype) and P3 (right; Bmp5^+/-^) mouse pup in which the pinna can be seen folding over itself at birth before unfolding and establishing its orientation.

Given the tissue level similarities between adult *Mus* and *Acomys* ear pinnae, we next sought to determine the extent to which ear pinna development in *Mus* mirrored that in *Acomys*. *Acomys* species are precocial and *A. cahirinus* have a gestational period twice as long as *Mus* (39-41 days vs. 20-21 days)(Figure 1E)(Brunjes, 1990; Haughton et al., 2016). Thus, a portion of ear pinna development in *Mus* that occurs postnatally, occurs neonatally in *Acomys* (Figure 1E). For example, prior to birth in *Mus*, the pinna was folded against the head before later unfolding distally from the skull at P3 (Figure 1F). In contrast, the ear pinna unfolds during embryonic development in *Acomys* (Figure 1F). Expectedly, P1 *Acomys* have hair follicles and more developed ear ridges. Together these observations suggest that development of the ear pinna is likely conserved among rodents, despite differences in maturity at birth.

### Hair follicle catagen precedes chondrocyte hypertrophy in the ear pinna

We next performed a longitudinal study of ear pinna development in outbred strains of *Mus* beginning at E20.5. Ear tissue was collected every other day up to postnatal day 21 (P21) to analyze development of the separate tissue types within the ear pinna (Figure 2, Supplemental Figure 2). At E20.5 the ear pinna was folded ventrally, and we observed mesenchymal cells aggregating in the center of the ear pinna (Figure 2A, S2). The pinna epidermis resembled a transitional epithelium where a stratum corneum was visible, but the stratum spinosum appeared poorly differentiated and the cells of the stratum basale lacked apical-basal polarity (Figure 2A). Interestingly, we observed interruptions in areas of central mesenchyme by lateral tissue extending medially (Figure 2B). At P1, condensing mesenchymal cells continued to form a dense aggregation while the dermal compartment of the pinna expanded resulting in decreased cellular density (Figure 2B). The epidermis now presented as a fully stratified epithelium, with a clearly differentiated stratum spinosum. Basal cells of the stratum basale were arranged in a columnar orientation and exhibited clear apical-basal polarity (Figure 2C). This transition of the epidermis was coincident with the onset of hair follicle morphogenesis and we observed hair germs beginning to form. Using an embryonic myosin heavy chain (Myh3) antibody we observed myofiber assembly along the dorsal surface of the future elastic cartilage (Figure 2D). P3 marked the onset of chondrogenesis as condensing mesenchymal cells began secreting Tenascin-C (Figure 2E). Hair follicle morphogenesis continued at P5 with extension of follicles into the dermis coincident with when the ear unfolds (Figure 2F, Supplemental Figure 2). Since muscle can be seen developing on the dorsal side of the ear pinna, contraction of this muscle as it matures may contribute to unfolding of the ear pinna. As chondrogenesis proceeded (P7-P9), Tenascin-C became localized to peripheral cells surrounding maturing chondrocytes and mesenchymal cells began depositing extracellular matrix within the dermis (Figure 2F-H). Adipocytes became visible at P9 (Figure 2I). A mature cartilage matrix, marked by the presence of glycosaminoglycan deposition, is absent at P7, but begins forming proximally at P9 (Fig 1J). Interestingly, while this maturation appears to be occurring proximal to distally, there are multiple niduses of maturation. At P11 mature blood vessels can be seen extending between the dorsal and ventral mesenchyme, interrupting the developing cartilage (Figure 1K). Hair follicle morphogenesis continued through the second week and sebaceous glands were visible by P15 accompanying the individual hair follicles (Figure 2L)(Muller-Rover et al., 2001; Paus et al., 1999). As the hair follicles completed catagen, chondrocytes continued maturing such that by P21 vacuolated cells were observed interspersed between chondrocytes and Tenascin-C marked the mature perichondrium (Figure 2L-N). At P21 the tissue architecture in *Mus* reflected the adult condition except for complete maturation of the elastic cartilage. Together our data shows that elastic cartilage chondrogenesis and hair follicle morphogenesis occur in parallel within the ear pinna and that differentiation of mature chondrocytes appeared to coincide when hair follicles complete catagen. This survey allows us to establish presumptive timing of development of the ear pinna tissues (Figure 2O).

**Figure 2.**
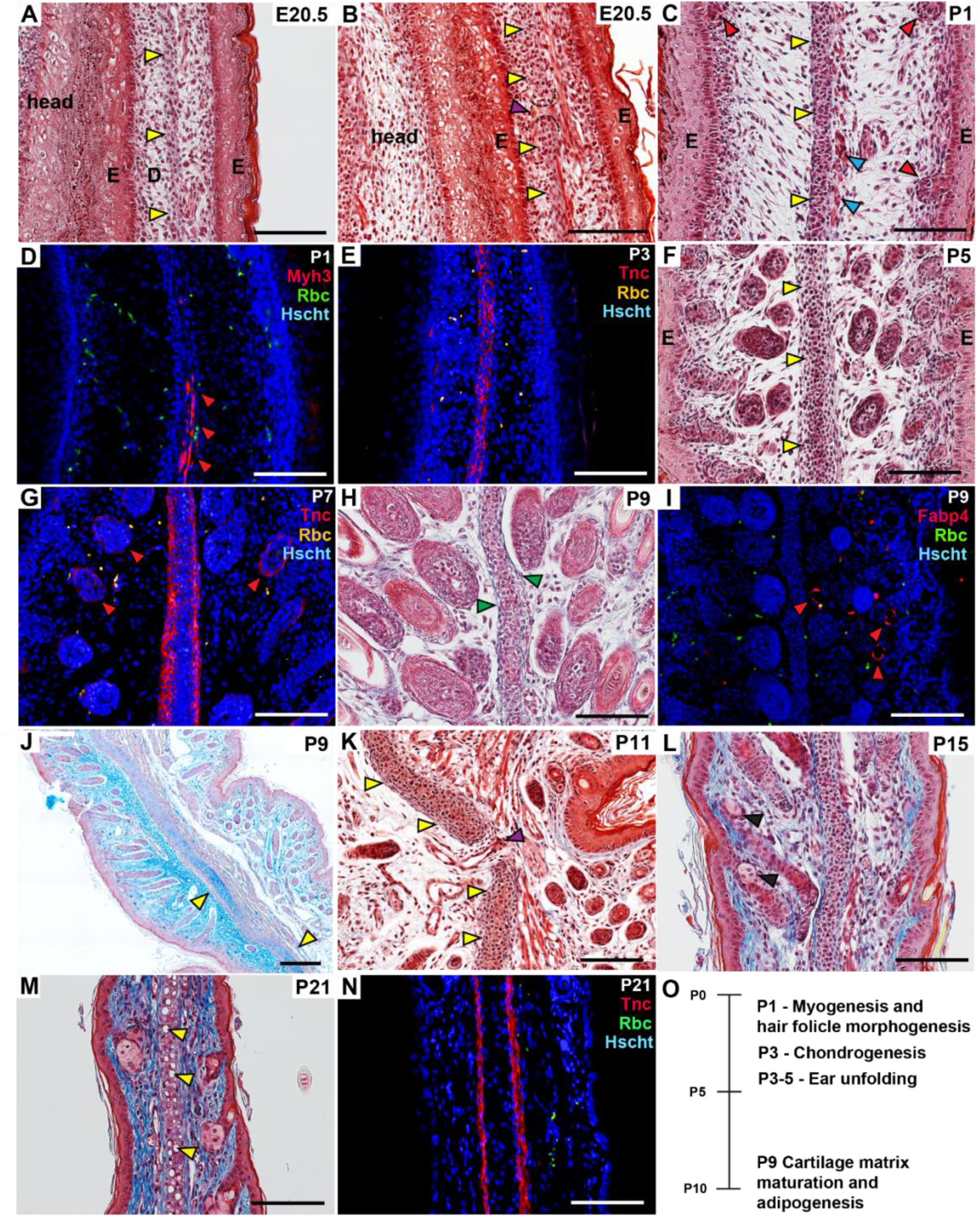
Ear pinna development in *M. musculus*. Masson’s Trichrome (A, B, C, F, H, K, L,M) staining, alcian blue (J) staining and immunostaining (D, E, G, I, N) of developing ear pinna in *Mus*. All sections are oriented with the dorsal side to the left and the proximal side to the bottom. In trichrome stained sections, nuclei are stained black, cytoplasm is stained pink/red and connective tissue and extracellular matrix are stained blue. Colored arrows indicate specific anatomical structures that arise at specific stages during development: cartilage (yellow) perichondrium (green), hair follicles (red), sebaceous glands (black) and presumptive and mature vasculature (purple). Dotted lines indicate interruption in central condensed mesenchyme (B). Positive immunostaining for Myosin Heavy Chain 3 (D), Tenascin-C (E, G, N) and Fabp4 (I) with red arrows indicating positively labeled cells. All scale bars are 100 µm unless specified otherwise. RBC = red blood cell. Hair follicle staging during morphogenesis was classified according to published staging guides (Muller-Rover et al., 2001; Paus et al., 1999). n=5 for E20.5, P1, P3, P5, P7, and P5 and n=3 for P9, P11, P15, and P21. (O) Approximate stage of *Mus* ear pinna developmental milestones based on histological survey.

### Developing Acomys perichondrium does not express Tenascin-C

We repeated our developmental analysis in *Acomys* using tissue that was collected at varying time points (roughly every 5 days) beginning at E20 until maturation (Figure 3; Supplemental Figure 3). The development of the primary tissue compartments (epidermis, hair follicles and cartilage) largely mirrored what was observed in *Mus*, with a few notable differences. First, due to precocial development in *Acomys*, pre-cartilaginous mesenchymal condensation occurred *in utero* as did the beginning phases of hair follicle morphogenesis (Figure 3A-F). Like *Mus*, early in development the *Acomys* ear is folded ventrally, however unfolding occurs in utero between E20 and E25 (Supplemental Figure 3). Of note, unfolding occurred prior to apparent hair follicle morphogenesis and chondrogenesis, making this the only developmental benchmark that occurs in an order different from *Mus*. Surprisingly, we could not detect Tenascin-C in condensing mesenchyme, despite a similar structure to *Mus* ears at the onset of Tenascin-C expression (Figure 3C). Mature vasculature can be seen interrupting the presumptive cartilage, likely representing the foramen we observed in adult ears (Figure 3D). At P1, deposits of extracellular matrix were visible just below the epidermis and sebaceous glands had already begun to form in association with developing hair follicles (Figure 3G). Although the perichondrium had fully formed by P5 (Figure 3H) we did not detect Tenascin-C at this stage or later (Figure 3K) despite the antibody recognizing Tenascin-C in this species. (Supplemental Figure 4). Despite these rather subtle differences, our analysis supports a general developmental scheme for the external ear across rodents. These data show that while the developmental stage at which the benchmarks of ear development occur vary among *Mus* and *Acomys*, the order and mechanism of these benchmarks is highly conserved (Figure 3L)

**Figure 3.**
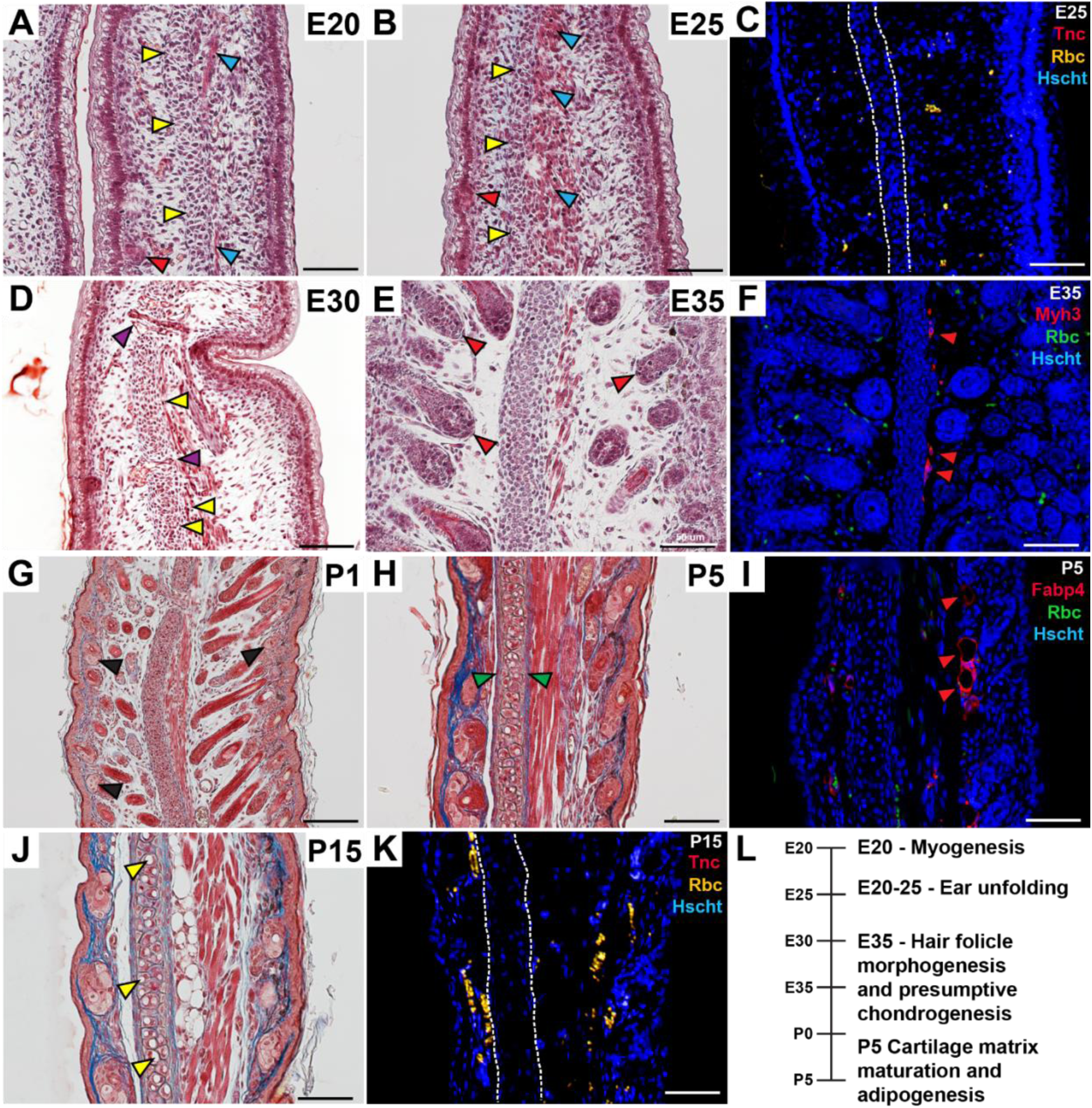
Ear pinna development in *A. cahirinus.* Masson’s Trichrome staining (A, B, D, E, G, H, J) and immunostaining (C, F, I, K) of developing ear pinna in *Acomys*. All sections are oriented with the dorsal side to the left and the proximal side to the bottom. In trichrome stained sections, nuclei are stained black, cytoplasm is stained pink/red and connective tissue and extracellular matrix are stained blue. Colored arrows indicate specific anatomical structures that arise at specific stages during development: muscle (cyan), cartilage (yellow), perichondrium (green), hair follicles (red), sebaceous glands (black), and vasculature (purple). Positive immunostaining for Tenascin-C (C, K), Myosin Heavy Chain 3 (F) and Fabp4 (I) with red with red arrows indicating positively labeled cells. Dashed lines indicate the boundaries of the condensing cells (C) and the perichondrium (K). All scale bars are 100 µm unless specified otherwise. RBC = red blood cell. n=1 for E20, E25 and E35 and n=2 for P1, P5 and P15. (L) Approximate stage of *Acomys* ear pinna developmental milestones based on histological survey.

### Wnt1 lineage tracing reveals that ear pinna dermis, adipose tissue and elastic cartilage are derived from neural crest cells

Previous work inactivating *hoxa2* in mice (completely or conditionally) demonstrated the importance of neural crest migration through the first pharyngeal arch for pinna formation (Rijli et al., 1993; Santagati et al., 2005). Examining mouse embryos up through E18.5, a recent study concluded that all tissues in the ear pinna were of neural crest origin except for the epidermis (Minoux et al., 2013). To test this hypothesis in mature tissue, we tracked neural crest cells (NCC) using a widely employed *Wnt1-cre* driver that labels pre-migratory NCC crossed to *ROSA^mT/mG^* females (Lewis et al., 2013; Muzumdar et al., 2007). In the *ROSA^mT/mG^*reporter strain all cells express membrane-bound tdTomato unless they undergo a Cre-mediated recombination event in which case they will lose tdTomato and instead express EGFP. Thus, by crossing *Wnt1-cre* males to *ROSA^mT/mG^*females all pre-migratory neural crest cells and their progeny would by EGFP+/tdTomato-(Figure 4A-J).

**Figure 4.**
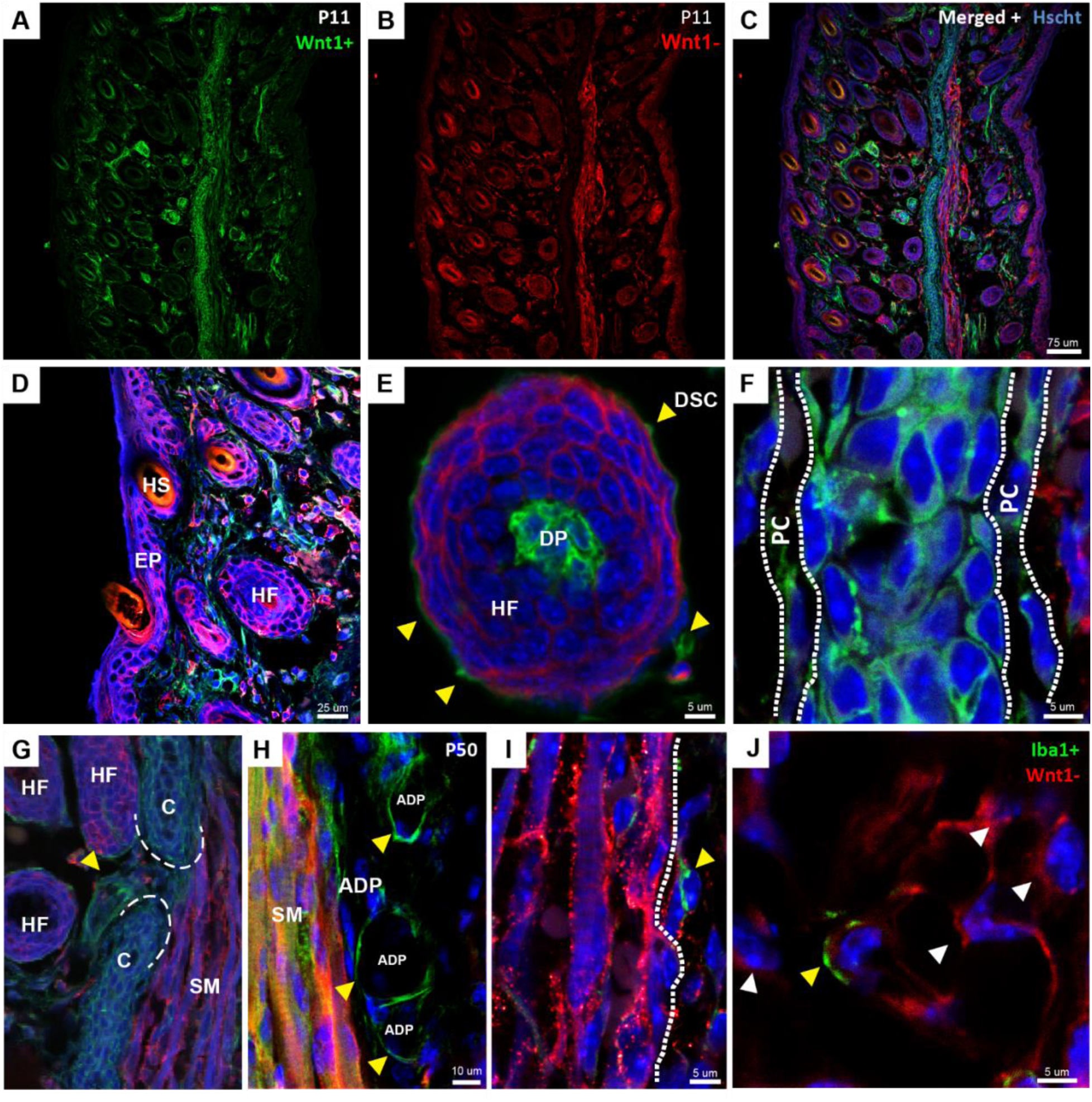
The ear pinna dermis, adipose tissue and elastic cartilage are derived from neural crest cells. Neural crest lineage tracing was performed by crossing male *Wnt1-Cre* driver mice to female *ROSA^mT/mG^* reporter mice (see Methods). Tissue was collected at P11 (all panels except H) and P50 (Panel H only). (A) *Wnt1+* structures are neural crest derivatives while (B) *Wnt1-* structures are not. (C) Overlay of *Wnt1+*, *Wnt1-* and Hoescht staining at P11. (D) High magnification of *Wnt1-* epidermis (EP). The hair shaft (HS) and hair follicle (HF) are also *Wnt1-*. (E) High magnification of a hair follicle, where the dermal papilla (DP) and dermal sheath cells (DSC, yellow arrows) are *Wnt1+* while the rest of the follicle is *Wnt1-*. (F) High magnification of the *Wnt1+* elastic cartilage and perichondrium (PC, white dashed line). (G) High magnification of cartilage foramen (white dashed lines) in which an apparent *Wnt1+* blood vessel runs (yellow arrow). (H) High magnification of *Wnt1+* adipocytes (ADP) marked by yellow arrows at P50. (I) High magnification of *Wnt1-* multinucleated muscle fibers with a *Wnt1+* cell (yellow arrow) flanking it. (J) High magnification of a macrophage marked by Iba1 (yellow arrow) within a population of *Wnt1-* cells (white arrows) within the dermis. Immunostaining for Iba1 was done using a far red (Cy5 Affinipure) secondary antibody to distinguish positively signal from endogenous tdTomato. n=3 for all panels.

Using this strategy, we first examined the ear pinna at P11 when all adult tissue structures in the ear pinna are present and observed Wnt1+ (GFP+) and Wnt1-(TOMATO+) cells (Figure 4A-C). Wnt1-cells were observed throughout the epidermis and in most of the hair follicle (Figure 4D-E). However, while most components of the hair follicle were Wnt1-, dermal papilla cells and cells in the dermal sheath were Wnt1+ (Figure 4E). In addition to dermal papilla and sheath cells, we observed Wnt1+ cells contributing completely to the elastic cartilage, perichondrium, dermis, and adipose tissue (Figure 4F-H and Supplementary Figure 5A-B). Within cartilage foramen we also identified Wnt1+ cells in what appeared to be vasculature, suggesting that blood vessels within the ear are also derived from cranial neural crest (Figure 4G). We then examined ear tissue at P50, well after development of the pinna was complete, but when more adipocytes were present (Figure 4G and Supplementary Figure 5C). To identify mature adipocytes, we used Fabp4 to double-label Wnt1+ cells which revealed a neural crest origin for adipocytes in the ear pinna (Figure 4G and Supplementary Figure 5C). We found that multi-nucleated muscle fibers comprising the skeletal muscle were not derived from neural crest (Figure 4I). In addition to muscle, we detected a small population of Wnt1-cells interspersed throughout the mesenchymal compartment. We tested if these cells were circulating or tissue-specific macrophages using a pan macrophage marker (Iba1) and detected a small number of Wnt1-/ Iba1+ cells (Figure 4J). It is also likely that some of the mesenchymal cells within the Wnt1-population (about 16%) are neural crest in origin but did not undergo successful Cre-recombination, as recombination using the *wnt1-cre* line is <100% (Hari et al., 2012). Taken together, our lineage analysis shows that neural crest cells give rise to elastic cartilage, perichondrium, dermis, vasculature, and adipose tissue, while skeletal muscle in the ear pinna derives from a non-neural crest source. In addition, head neural crest cells give rise to the dermal papilla and dermal sheath cells of the ear pinna hair follicles. These results are consistent with the contribution of cranial neural crest cells to other tissues of the head and provide a detailed developmental origin for the cell types of the ear pinna(Roth et al., 2021).

### Bmp5 is critical for elastic cartilage maturation

To extend our cellular analysis of ear pinna development, and better understand the process of auricular elastic cartilage maturation, we sought to investigate the underlying defect that produces deformed ear pinnae in the naturally occurring *short ear* mouse strain (Green and Green, 1942). *Short ear* (*bmp5^se/se^*, referred to henceforth as Bmp5^-/-^) mice have a C to G point mutation in the final exon of *bmp5* which results in a premature stop codon and causes skeletal defects throughout the body(King et al., 1994). Among these skeletal defects, *short ear* mice derive their name from a truncated ear pinna (Green and Green, 1942; Lynch, 1921). Green and Green (1942) performed a histological analysis of Bmp5^-/-^ animals at early post-natal time points P1, P3 and P6 and showed that condensing elastic cartilage in the outer pinna (scapha) was more diffuse than wild-type ears as early as P3 while no other tissue compartments appeared to be affected. They also noted that ears in P14 wild-type animals begin to elongate whereas this elongation was truncated in Bmp5^-/-^ animals(Green and Green, 1942). Building on this work, we compared Bmp5^-/+^, whose ears develop normally with Bmp5^-/-^ adult mice and observed that the scapha appeared furrowed and truncated in Bmp5^-/-^ mice (Figure 5A and B). We next examined the cellular architecture of ear pinnae from Bmp5^-/+^ and Bmp5^-/-^ mice (Figure 5C-H). Based on the *bmp5* mutation we predicted that ears from Bmp5^-/-^ mice would have elastic cartilage defects throughout the pinna. Surprisingly, we found that the proximal portion of the ear pinna exhibited relatively normal tissue architecture including a well-developed elastic cartilage plate and associated skeletal muscle (Figure 5D). In addition, the most distal tip of the ear pinna appears relatively unperturbed in *short ear* mutants. However, examining the truncated regions of the ear pinna (specifically the scapha) we found disorganized cartilage nodules instead of a continuous plate and increased abundance of adipose cells. Examination of developing Bmp5^+/-^ and Bmp5^-/-^ ears shows the stages of development mirrored wildtype *Mus* (Supplemental Figure 7). Bmp5^+/-^ and Bmp5^-/-^ ears are histologically indistinguishable at early time points (P1-P3), with a robust condensation of mesenchymal cells running down the center of the ear. But starting at P5 this condensation appeared to narrow in Bmp5^-/-^ ears with cartilage failing to develop normally in the distal ear. Using pSMAD 1/5/8 as a downstream readout of Bmp-signaling, we saw pSMAD 1/5/8+ cells within the condensing mesenchyme through P1 in wildtype mice (Supplementary Figure 6A-B). However, at P3, pSMAD 1/5/8+ cells were no longer present in this condensing population, suggesting that *Bmp5*^-/-^ defects may be established as early as P1, despite no observable histological changes (Supplementary Figure 6C).

**Figure 5.**
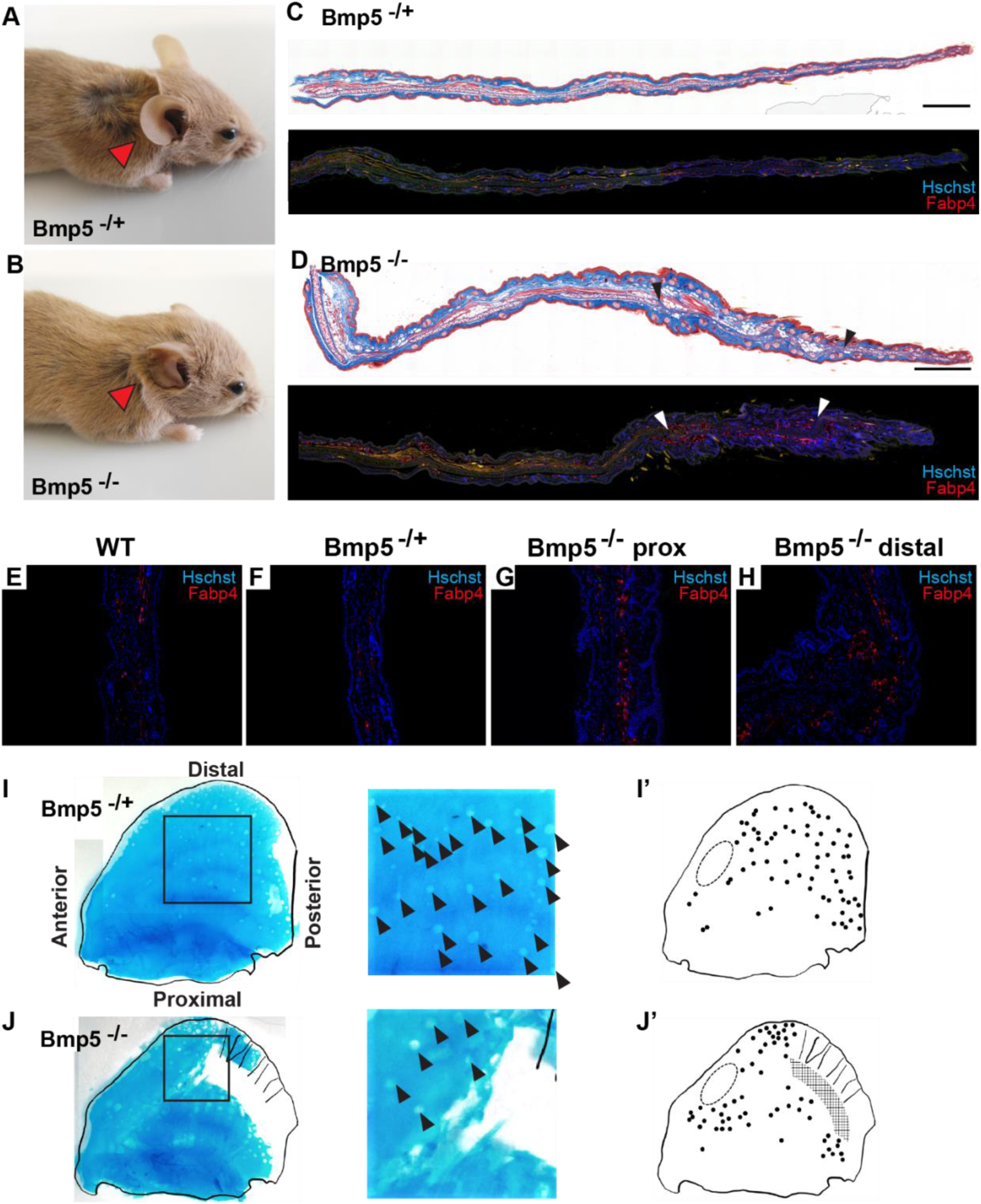
Bmp-signaling is critical for elastic cartilage development in the ear pinna. (A-B) Differences in ear pinna size and shape in Bmp5 heterozygous (A) and null (B) mice can be observed. (C-H) Histological comparison of Bmp5^+/-^ and Bmp5^-/-^ null mice using Masson’s Trichrome (C and D) and Fabp4 immunostaining (C-H). Regions of patterned and mispatterned elastic cartilage and increased adipose abundance are observed within the distal pinnae of null mice, black and white arrowheads (D). n=3 for each group. (I-J) Whole mount alcian blue staining of adult Bmp5^+/-^ (E) and Bmp5^-/-^ (F) mice ear pinna elastic cartilage. Ear outline generated from partially intact ear (see Supplemental Figure 1). Arrowheads indicate foramen within the cartilage structure. (E’-F’) Schematic representation of foramen visible in the adult *Mus* (E’) and Acomys (F’) ear pinna elastic cartilage. Dotted oval indicates area of uninterrupted cartilage at the anterior edge. Crosshatch indicates discontinuous cartilage area in Bmp5^-/-^ ear. n=2 for Bmp5^+/-^ *Mus* and n=1 for Bmp5^-/-^ *Mus* alcian blue whole mount stains.

In order, to further analyze this apparent tradeoff between chondrogenesis and adipogenesis, we used Fabp4 (fatty acid binding protein), a marker of mature adipocytes, to compare the proportion of adipocytes in wild-type, *Bmp5*^+/-^ and in the proximal and distal compartments of Bmp5^-/-^ mice (e.g., Bmp5^-/-^ prox and Bmp5^-/-^ distal) (Figure 5E-H). Using this approach we found that both the Bmp5^+/-^ and Bmp5^-/-^ mice had significantly more adipocytes within the dermis than wild-type (One-way ANOVA, Tukey’s, Bmp5^+/-^: 13.07 ± 1.52% and Bmp5^-/-^: 15.34 ± 1.17% versus wild-type: 7.45 ± 0.86%, p < 0.05). Finally, using whole mount alcian blue staining we examined the gross cartilage structures of Bmp5^+/-^ and Bmp5^-/-^ adult mice. We found that *Bmp5*^+/-^ mice have a continuous cartilage sheet like other mouse strains, with foramina scattered more densely in the distal and caudal quadrants as well as a dense area of cartilage along the rostral edge (Figure 5I-J’). Interestingly, Bmp5^-/-^ mutants also had an uninterrupted area of cartilage at the rostral edge, surrounded by foramina (Figure 5J-J’). However, cartilage in the caudal half of the ear was discontinuous, making it impossible to recover using this method. This area of discontinuous cartilage coincided with those areas with the most apparent loss of structure in our histological sections. These results using Bmp5^-/-^ mice suggest that mice deficient in both *bmp5* allele(s) may have aberrations in neural crest cell differentiation along the chondrogenic and adipogenic lineages.

### Bmp5^-/-^ Mus fail to develop mid-distal ear cartilage

To more completely understand the elastic cartilage defects occurring in Bmp5^-/-^ mice, we next examined chondroprogenitor differentiation and perichondrium development in Bmp5^+/-^ and Bmp5^-/-^ mice. At P5, chondroprogenitors marked by Sox9 expression were much denser in the medial condensation of presumptive cartilage in Bmp5^+/-^ ears than Bmp5^-/-^ ears (Figure 6A). Sox9 also marked dermal papilla and sebaceous glands, which appeared at similar densities in these two mouse types. In the center of the ear at P7, we observed two distinct sheets of cartilage in Bmp5^+/-^ ears interrupted by a small foramen (Figure 6B). While there were also two areas of cartilage observable at the proximal and distal ends in Bmp5^-/-^ ears, they were separated by a large gap (Figure 6B’). We examined perichondrium development in P7 ears and found that TNC nicely outlined condensations of presumptive cartilage in Bmp5^+/-^ ears (Figure 6C). In mutant ears within the Sox9-central area, TNC+ cells appeared compressed suggesting that the perichondrium had successfully formed, but cartilage differentiation had stalled (Figure 6C’). This observation supports that perichondrium development across the ear does not require cartilage development. We measured the distance between the end of the most proximal piece of cartilage to the end of the ear (C1 to ear end) and found this length was not significantly different between Bmp5^+/-^ and Bmp5^-/-^ ears (Figure 6D and E). However, there was a significant increase in the distance between the end of the proximal cartilage sheet and the distal cartilage sheet (C1 to C2) and a significant decrease in the length of the distal sheet (C2 to ear end) in Bmp5^-/-^ compared to Bmp5^+/-^. These data suggest that the location of the foramen may correspond to breaks in independently maturing areas of chondrogenesis.

**Figure 6:**
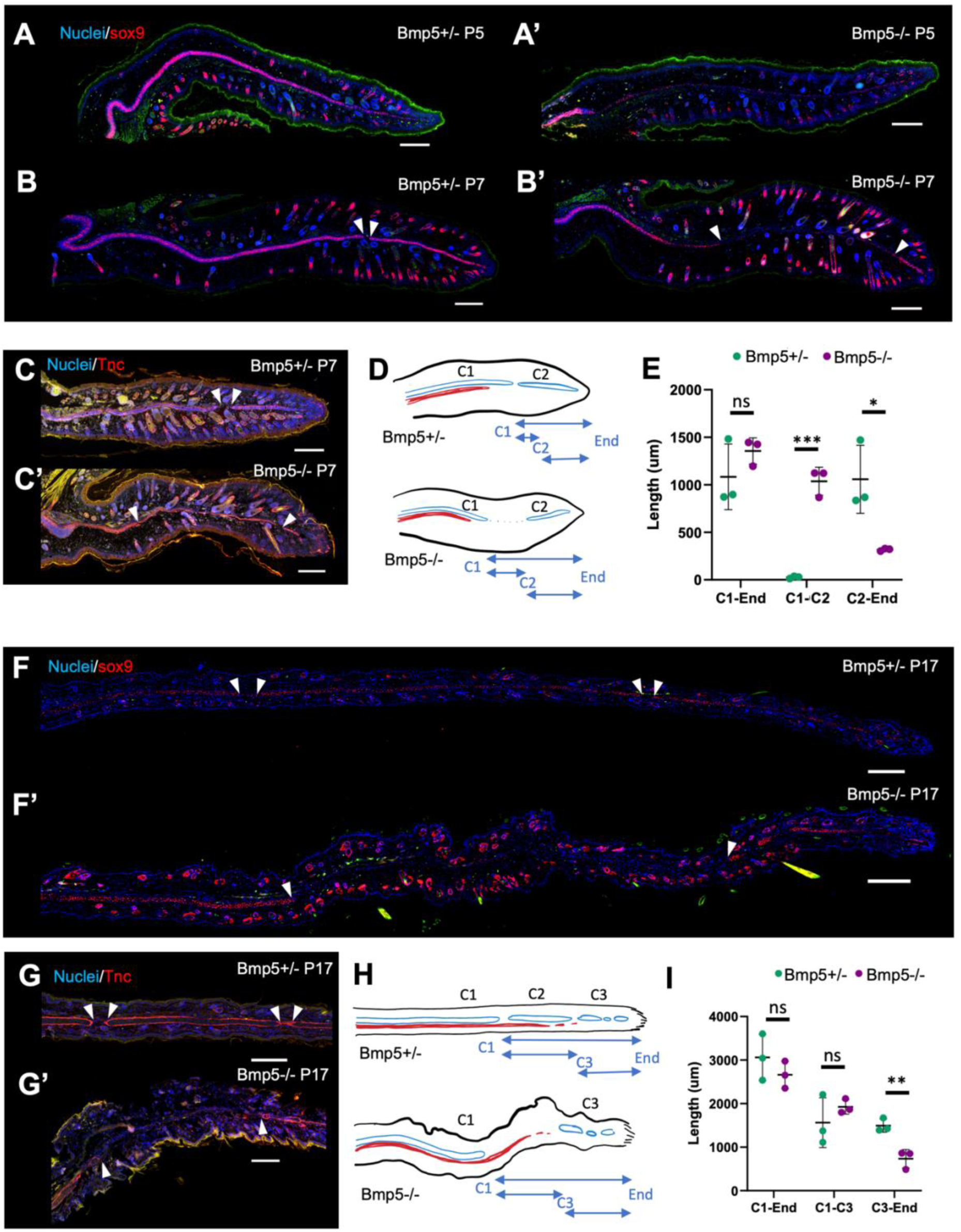
Bmp5^-/-^ mice fail to develop cartilage in the mid distal ear pinna. (A-C, F, G) Histological comparison of Bmp5^+/-^ and Bmp5^-/-^ mice using Sox9 (A, B, and F) and Tenascin (C and G) immunostaining. White arrowheads indicate proximal or distal limits of continuous cartilage sheets separated by foramina. (D) Schematic representation of developing cartilage in P7 Bmp5^+/-^ and Bmp5^-/-^ null mice. Blue arrows indicate measurements compared in (E). (H) Schematic representation of developing cartilage in P17 Bmp5^+/-^ and Bmp5^-/-^ mice. Blue arrows indicate measurements compared in (I). n=3 for all images. (* P<0.05, ** P<0.01, *** P<0.001).

At P17, three distinct sheets of cartilage were observable across the Bmp5^+/-^ ear, while in the Bmp5^-/-^ ear there were only two areas of Sox9 expression with a large gap in between (Figure 6F). This organization was also visible when staining the perichondrium with TNC (Figure 6G). which revealed a gap in the mutant suggesting chondrocytes are necessary for maintenance of the perichondrium. Intriguingly, the distance between the ends of the most proximal (C1) and distal (C3) cartilage condensations in Bmp5^+/-^ ears is about the same length as the Sox9-C1 to C3 gap in Bmp5^-/-^ ears (Figure 6H and I). While the separation between these sheets represent foramen, rather than a continuous break in cartilage across the ear, they still may approximately mark the boundaries of three separate areas of cartilage development in the rodent ear. In the case of Bmp5^-/-^ ears, the middle condensation (C2) fails to develop into chondroprogenitors and later mature cartilage. Interestingly, in Bmp5^-/-^ P7 and P17 ears, muscle appears to develop normally, extending into the mid ear even without the presence of a cartilage scaffold (Supplemental Fig. 8). Thus, the perichondrium may be sufficient to guide muscle development in Bmp5-/-mice, or muscle development may occur independently of cartilage development. These observations provide further insight into how elastic cartilage of the ear pinna develops and how it interacts with the developing perichondrium and muscle.

### Chondroprogenitors in Bmp5^-/-^ ears fail to expand through proliferation

To determine if the loss of Sox9+ cells in the developing Bmp5*^-/-^* mutant ear resulted from developmental failure or misspecification, we looked at proliferation and adipogenesis in Bmp5^+/-^ and Bmp5^+/-^ ears. Previous studies suggested the presence of adipochondrocyte progenitors within the ear that can give rise to both cartilage and fat lineages(Sanzone and Reith, 1976). Considering the increase in adipocyte abundance we observed in adult mutant ears, we investigated if chondroblast differentiation was shifted toward adipogenesis.

We first marked cycling cells using Ki67, which is expressed by cells in S, G2, and M phase. While we did not observe significant differences among Ki67+ cells in the far proximal and distal presumptive cartilage condensations, there were significantly fewer Ki67+ cells in the mid-pinna condensation in Bmp5^-/-^ ears compared to Bmp5^+/-^ (Figure 7A and B). Proliferation within the mesenchyme, hair follicles and epidermis appeared unperturbed in both genotypes. These data suggest that chondroprogenitors fail to proliferate in the mid ear.

**Figure 7:**
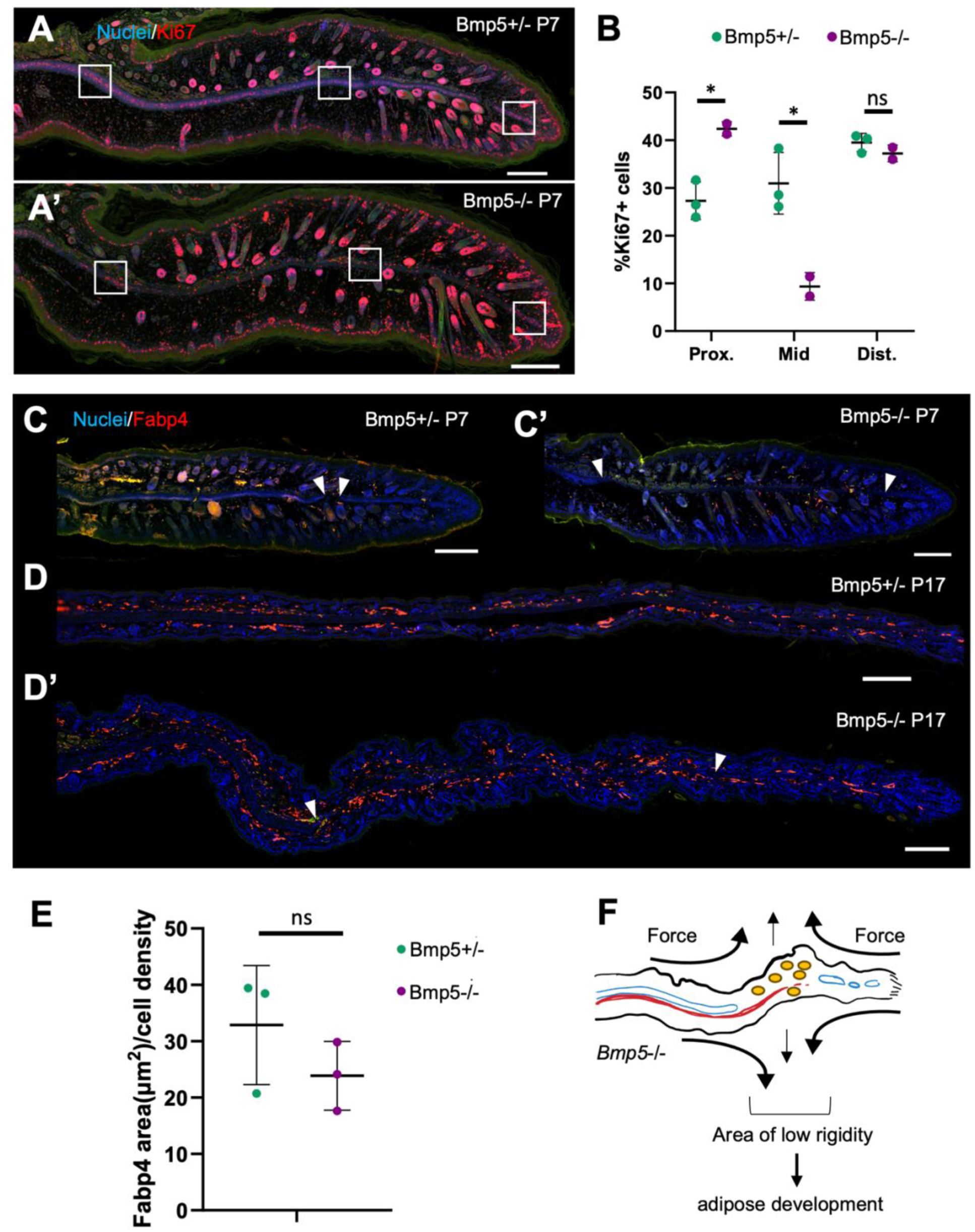
Bmp5^-/-^ ear pinna chondroprogenitors are deficient in proliferation, not differentiation. (A, C, D) Histological comparison of Bmp5^+/-^ and Bmp5^-/-^ mice using Ki67 (A) and Fabp4 (C and D) immunostaining. (B) Comparison of Ki67+ cells within the presumptive cartilage condensation of the proximal, mid, and far distal (white squares) ear pinna in P7 Bmp5^+/-^ and Bmp5^-/-^ mice. (E) Comparison of Fabp4+ area normalized to cell count in the mid-distal ear pinna in P17 Bmp5^+/-^ and Bmp5^-/-^ mice. (F) Hypothesized mechanical basis for increased adipogenesis in adult Bmp5^-/-^ mid-distal ear pinna. n=3 for all images. (* P<0.05).

Next, we used Fabp4 to examine the abundance of adipoprogenitors and adipocytes at P7 and P17. Interestingly, despite having more apparent fat in the adult ear, there did not appear to be any difference between Bmp5^+/-^ and Bmp5^-/-^ Fabp4+ cell abundance at P7 or P17 (Figure 7C-D). We measured the total area of Fapb4+ staining normalized to cell number in the mid ear and found there was no significant difference between Bmp5^+/-^ and Bmp5^-/-^ ears at P17 (Figure 7E). These data suggested that adipose tissue develops normally in Bmp5^-/-^ and the increased abundance of adipocytes observed in adults occurs post-developmentally. We hypothesize that the lack of mid ear cartilage in the Bmp5^-/-^ mutants results in a combination of a softer tissue environment and an increase in lateral forces by the adjacent tissue containing rigid cartilage. This combination of altered stiffness and de-compression forces may create a pro-adipogenic mechanical environment in adulthood (Figure 7F)(Hossain et al., 2010; Li et al., 2013; Su et al., 2022).

### Bmp5*^+/-^* Mus have impaired chondrogenesis in adulthood

To determine if Bmp5 is required for chondrogenesis in adulthood. We next challenged Bmp5^+/-^ and Bmp5^-/-^ mice with a full thickness injury to the ear pinna. Normally, *Mus* respond to these injuries with fibrosis, however within the scar they can form new cartilage nodules, showing they are still able to activate chondrogenesis in the adult ear (Gawriluk et al., 2016; Leung et al., 2015). We created 4mm hole punches in the center of Bmp5^+/-^ and Bmp5^-/-^ ears allowing us to look at chondrogenesis adjacent to the proximal cartilage, and with two adjacent 2mm punches through the distal ear to observe chondrogenesis adjacent to the mid-distal ear cartilage (Figure 8A). This specifically allowed us to assess injury responses in the normal and abnormal cartilage areas of Bmp5^-/-^ mice. We measured closure of the wound over 50 days and compared them to our outbred ND4 strain which generates robust scar tissue in response to wounding. Interestingly, we found that 4mm holes closed similarly in wildtype and Bmp5^-/-^ mice, while Bmp5^+/-^ closed significantly less tissue (2-Way ANOVA, analyzing genotype and time as variables, Bmp5^+/-^ vs WT or Bmp5^-/-^, genotype factor was significantly different (P<0.0001), WT vs Bmp5^-/-^, genotype factor was not significantly different (P=0.5111)) (Figure 8B). Because we could not ensure consistency of the original wound size in distal 2mm ear punches we could not statistically assess closure of these injuries. However, because Bmp5^-/-^ had less rigidity to their ears we found they were more prone to deformation of the hole (Supplemental Figure 9).

**Figure 8:**
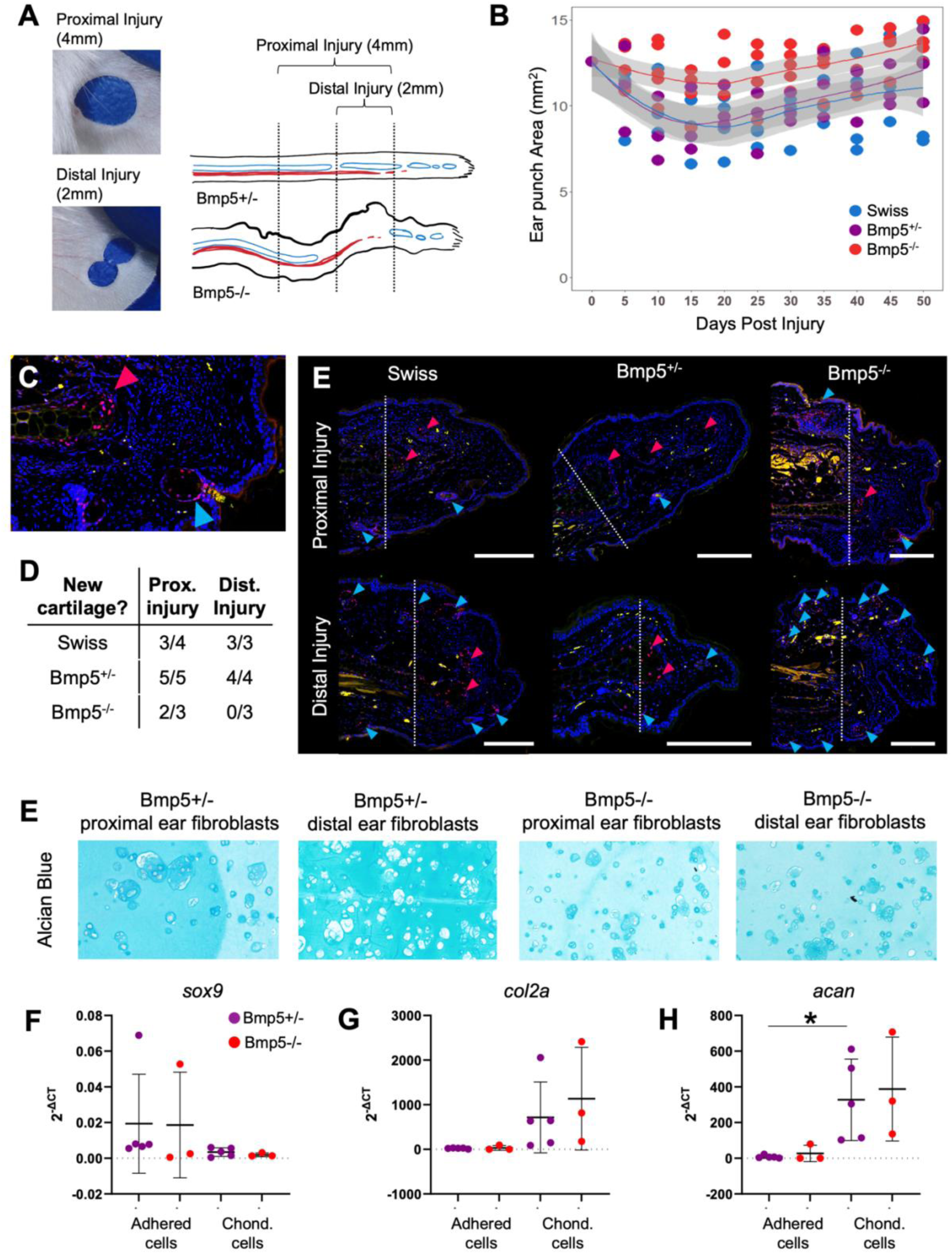
Bmp5-/- *Mus* have impaired chondrogenesis following wounding in the distal ear pinna. (A) Representative images and schematic representation of two different wounding paradigms: A 4mm ear punch in which the proximal wound edge passes through intact cartilage of both Bmp5+/- and Bmp5-/- mice, and a 2mm distal ear punch in which the proximal wound edge passes through the mid-distal ear, which has discontinuous cartilage development in Bmp5-/- mice. (B) Ear punch area over time following proximal 4mm full thickness ear punch in wildtype (Swiss webster; blue), Bmp5+/- (purple), and Bmp5-/- (red) mice. Each point represents an individual mouse. Lines represent average ear hole area with 95% confidence area highlighted in gray. (C) Table showing proportion of mice of each genotype that had new cartilage formed following a proximal 4mm full thickness ear punch or distal 2mm full thickness ear punch. (D) High magnification image showing *Mus* ear pinna scar tissue containing new cartilage (red arrowhead) and hair follicles/glands (blue arrowhead), both of which stain positive for Sox9. New cartilage is distinguishable by its structure and central location, while hair follicles/glands are found on the periphery. (E) Histological comparison of scars in different mouse genotypes following proximal 4mm or distal 2mm injury using sox9 immunostaining. Red arrowheads indicate new cartilage nodules, blue arrowheads indicate hair follicles and glands. Scale bars = 100um. (F) Alcian blue stain of 3D cell cultures of Bmp5+/- and -/- fibroblasts isolated from the proximal or the distal ear pinna following in vitro chondrogenesis. Pictures were selected from cells with mature cartilage marker expression closest to the mean based on QPCR analysis (see next panel). (G-I) QPCR expression analysis of *sox9* (G), *collagen2a* (H), and *aggrecan* (I) in Bmp5+/- and -/- fibroblasts before and after chondrogenesis. (* P<0.05).

We also assessed chondrogenesis in scar tissue using Sox9 staining 85 days post injury. This marker stains both chondroprogenitors and hair follicles which can be differentiated from each other based on morphology and location (Figure 8C). We found that in all three genotypes the 4mm proximal wound resulted in new cartilage formation within the scars of most ears (Figure 8D and E). Interestingly, while wildtype and Bmp5^+/-^ mice were able to generate cartilage in distal wounds, none of the Bmp^-/-^ mice were able to generate new Sox9+ cartilage in their distal wounds. Notably, unlike in wildtype and Bmp^+/-^ ears, distally wounded Bmp^-/-^ did not have increased Sox9+ staining at the cartilage end to indicate increased activation of chondroprogenitors. Positive staining of hair follicle stem cells demonstrated that the Sox9 stain was working. Together, our data showed that Bmp^-/-^ mice continue to have impaired chondrogenesis in adulthood. However, this impairment is spatially limited to areas in which cartilage formation failed during development.

Lastly, to test if dermal fibroblasts from heterozygous and mutant mice could contribute to chondrogenesis we used an *in vitro* chondrogenesis assay for fibroblast differentiation potential. Using this assay, we found that fibroblasts from both genotypes were able to generate a glycosaminoglycan rich matrix in chondrogenic conditions (Figure 8F). QPCR for *sox9*, which marks early chondrogenesis, did not show increased expression in chondroid cells, likely because it is downregulated during maturation (Figure 8F). QPCR for markers of mature cartilage showed that both Bmp^+/-^ and Bmp^-/-^ fibroblasts that underwent chondrogenesis trended toward upregulated expression of *collagen 2a* and *aggrecan* (Figure 8G and H). These data show that Bmp5 is not required for fibroblasts to undergo chondrogenesis *in vitro*. Since our Bmp^-/-^ mice were not able to generate new cartilage after wounding, these data support that despite the capacity of fibroblasts to form new cartilage, only chondroblasts from existing cartilage give rise to new cartilage in the scar.

## DISCUSSION

Here we set out to study external ear pinna development in murid rodents and established a conserved sequence of events that leads to the coordinated formation of mature skin, elastic cartilage, skeletal muscle and adipose tissue architecture in the pinna. Through lineage tracing, we demonstrate that connective tissue fibroblasts, elastic cartilage, dermal papilla cells, dermal sheath cells, vasculature, and adipocytes in the adult pinna are derived from cranial neural crest. Although this study only included two murid species, *Acomys* and *Mus*, the developmental sequence presented in this study likely represents the general pattern for all rodents. Using *short ear* mutant mice harboring a point mutation in *bmp5*, we show that *bmp5* is exclusively necessary for development of the middle and part of the distal cartilage of the scapha. Our collective findings have implications for future development and regeneration studies of the ear pinna including congenital deformities of the ear pinna such as microtia and anotia.

The analgen to the pinna forms as an outgrowth from side of the head above the external acoustic meatus (EAM) and has been well described in mice up through E18.5 (Fuchs and Tucker, 2015; Minoux et al., 2013). Based on our data from embryonic stages, cells were visible at high density within the central region of the developing pinna and we found these cells were the precursor to elastic cartilage. Intriguingly at this stage, well before chondrogenesis ensues, lateral mesenchymal tissue can be seen extending between the rostral and caudal sides of the ear interrupting high density condensations. These interruptions are later observed to contain vasculature, suggesting that development of the vascular system is coincident with the development of the ear pinna cartilaginous structure. As the centrally located cells underwent chondrogenesis, the dermal compartment expanded in parallel with hair follicle and muscle morphogenesis. Following hair follicle catagen, maturation of the elastic chondrocytes continued to give rise to the mature cartilage plate.

One interesting difference we observed in this conserved developmental sequence concerned the expression of Tenascin-C. In *Mus*, we observed Tenascin-C expression beginning at P3 as Bmp-signaling in these same cells was downregulated. As chondrogenesis proceeded, Tenascin-C became excluded from the most central cells and was retained by peripheral cells. Ultimately, Tenascin-C expression became restricted to the perichondrium where it persisted in adults. In contrast, we were unable to detect Tenascin-C expression at any time during chondrogenesis in *Acomys* or in mature *Acomys* perichondrial cells even though it is upregulated during ear pinna regeneration (Gawriluk et al., 2016). This finding suggests that perichondrial cells may possess inherent differences across rodent species that will require further examination. Despite this interesting difference, our finding that *Mus* and *Acomys* largely share this general developmental plan allow us to conclude that *Mus* is a useful model to investigate ear pinna development as a proxy for *Acomys* and other murids. Recently, *Acomys* have emerged as a bonafide mammalian model of epimorphic regeneration. Compared to all other murid rodents studied to date who heal injuries via fibrotic repair, spiny mice are capable of re-growing full-thickness body skin and musculoskeletal tissue in the ear pinna (Gawriluk et al., 2016; Matias Santos et al., 2016; Seifert et al., 2012; Simkin et al., 2017). Our data provide a firm developmental context for future cellular studies of musculoskeletal regeneration using the ear pinna model which as part of the craniofacial complex has a unique developmental trajectory compared to body tissues.

Previous work showed that *rhombomere4-* and *Hoxa2*-expressing neural crest cells (NCC) of the second pharyngeal arch give rise to the embryonic pinna and that *Hoxa2* controls pinna morphogenesis via regulation of *Bmp4* (binding upstream of transcription start site) and *Bmp5* (binding in the third exon) (Minoux et al., 2013). While this work demonstrated that origin of the early pinna was neural crest (NC) derived, we sought to extend this work and ask whether tissues of the adult pinna were in fact all NC-derived or if non-NCCs migrated into the ear during later stages of pinna morphogenesis. Using lineage analysis of NCCs in mice we show that NC cells are the source of elastic cartilage, perichondrium, dermis, vasculature, and connective tissue of the ear, whereas the muscle is derived from a different germ layer (likely mesoderm). Whether muscle precursors migrate simultaneously or subsequent to cranial neural crest cells will require further investigation in the future.

In mice with *hoxa2* deletions, the expression of both Bmp4 and Bmp5 is significantly downregulated (Minoux et al., 2013). This work showed that Bmp4 and Bmp5 expression in the developing pinna were complementary, with Bmp4 primarily expressed primarily at the tip of the pinna and within a small percentage of the mesenchymal cells, while Bmp5 was expressed broadly throughout the mesenchyme at the base (Minoux et al., 2013). To explore how dysregulation of Bmp-signaling might affect the adult ear pinna we utilized a well-characterized mutant mouse strain, the *se* mouse, which carries a null mutation in *bmp5* and exhibits a number of skeletal defects (primarily in anatomical sites of elastic cartilage) including a truncated ear pinna(DiLeone et al., 2000; DiLeone et al., 1998; Klockars and Rautio, 2009). Interestingly, we found that tissue architecture was relatively normal in the proximal pinna (close to the head) and that cartilage defects manifested in the distal half of the pinna (scapha). In adult ears, the defected regions were void of elastic cartilage and accompanied by excess lipid deposition and an abundance of adipocytes within the dermis. Even in places where the cartilage was intact, we found significantly more adipocytes relative to wild-type mice. Interestingly, we saw also that Bmp5^-/+^ mice, whose pinnae exhibit normal tissue architecture, also have a significantly larger adipocyte population. It is likely that some degree of functional redundancy between Bmp4 and Bmp5 prevents the complete loss of elastic cartilage, although this hypothesis awaits further testing (Minoux et al., 2013). Interestingly, along the midline of the ear the boundaries of missing cartilage in Bmp5^-/-^ ears, corresponded to the location or foramen that form in the developing cartilage to allow passage of vasculature. Within this area of missing cartilage, chondroprogenitors fail to proliferate, though perichondrium still appears to develop. Cartilage proximal and distal to this zone of sparse proliferation develop normally, suggesting that even though the mesenchymal sheet that will eventually give rise to the cartilage appears to form simultaneously, there are multiple sites of chondrogenesis throughout this sheet. The consistency in the area of cartilage mis-patterned in the Bmp5^-/-^ mouse suggests a coordination of multiple well-demarcated sites of chondrogenesis within the ear. Further, our data show that *Bmp5* is exclusively necessary for the development of the mid-distal ear pinna cartilage.

While early histological studies have alluded to a specific adipochondrocyte progenitor cell, there is little molecular evidence from *in vivo* studies (Sanzone and Reith, 1976). However, *in vitro* stem cell differentiation assays have shown that mesenchymal stem cells derived from adipose, synovial membrane, bone marrow and other sources are multipotent and can be driven towards both an adipocyte and chondrocyte fate under specific culture conditions (De Bari et al., 2001; Dennis et al., 1999; Gimble and Guilak, 2003; Noth et al., 2002). We found that there was no increase in adipoprogenitors during ear pinna development of Bmp5^-/-^ mice. Coupled with our observation that Bmp5^-/-^ chondroprogenitors fail to proliferate, these data suggest that chondroprogenitors fail to establish in the ear scapha, rather than mis-differentiation along a different lineage. We hypothesize that increased adipocyte abundance in the ear scapha results from changes in tissue stiffness and directional forces due to loss of the cartilage scaffold. Adipogenesis is enhanced by soft substrates and inhibited by compressive forces. Loss of cartilage in that area likely creates an area of low stiffness and decreased compression (Hossain et al., 2010; Li et al., 2013; Su et al., 2022). The Bmp5^-/-^ mouse as a result may be an intriguing model for future studies into the biomechanical control of cell identity.

Finally, we show that impaired chondrogenesis within the ear scapha is retained in adulthood in Bmp5^-/-^ mice. When challenged with a full thickness ear punch wound, Bmp5^-/-^ mice can generate cartilage in scar tissue formed in the proximal ear, but not in the distal ear. Interestingly, dermal fibroblasts can undergo *in vitro* chondrogenesis even in the absence of Bmp5 (Culbert et al., 2014; Dastagir et al., 2014). However, the lack of cartilage formation in the distal wounds of Bmp5^-/-^ mouse ears suggests that the fibroblast lineage does not participate in chondrogenesis *in vivo*. Notably, we consistently see activation and accumulation of Sox9+ chondroprogenitors at the cartilage end of the wound boundary except in the distal ear of Bmp5^-/-^ mice. However, in Bmp5^-/-^ mice Sox9+ progenitors are still visible in the edges of the cartilage where they normally reside in the unwounded state. Our results suggest that Bmp5 may be required for activation of chondroprogenitors in the distal ear and these cells are exclusively required for chondrogenesis within scar tissue. Understanding the origin of the cells that give rise to the pinna has significant implications for regeneration, as it has been shown that in multiple vertebrate systems that the cells which replace the damaged tissue exhibit some degree of lineage restriction recapitulating their developmental origins (Kragl et al., 2009; Rinkevich et al., 2011; Singh et al., 2012). Cranial neural crest derived skin in humans has been found to heal faster and with less fibrosis compared to trunk skin, suggesting neural crest derived tissue has unique wound healing properties(Usansky et al., 2021). Interestingly, in our previous and ongoing regeneration studies, we notice a weaker regenerative potential in the ear skeletal muscle, a fact that may relate to the unique origin of this tissue compared to the rest of the pinna.

Together, our data establish a developmental time course and origin for the tissues of the ear pinna. In addition, they show differential signaling requirements in different parts of the ear pinna elastic cartilage that are retained into adulthood. These data contribute to our broader understanding of cranial neural crest tissue development and establish a basis for future wound healing and bioengineering strategies.

## ACKNOWLEDGMENTS

We thank all members of the Seifert lab for insightful discussions and Adam Cook and Joshua Sarli for animal husbandry. This work was supported by grants from the National Science Foundation (NSF) and the Office for International Science and Engineering (OISE) (IOS-1353713) and from the National Institute of Musculoskeletal, Arthritis and Skin Diseases (NIAMS) (R01AR070313) to A.W.S. The content in this article is solely the responsibility of the authors and does not necessarily represent the official views of the National Institutes of Health.

## AUTHOR CONTRIBUTIONS

RSA, SKB and AWS designed the project and experiments. RSA, SKB performed all the experiments with help from CKH. RSA, SKB, CKH and AWS analyzed the data and wrote the manuscript. All authors commented on and edited the final version.

