## Supplemental figs for "Neural crest cells give rise to non-myogenic mesenchymal tissue in the adult murid ear pinna"

**Supplemental Figures**

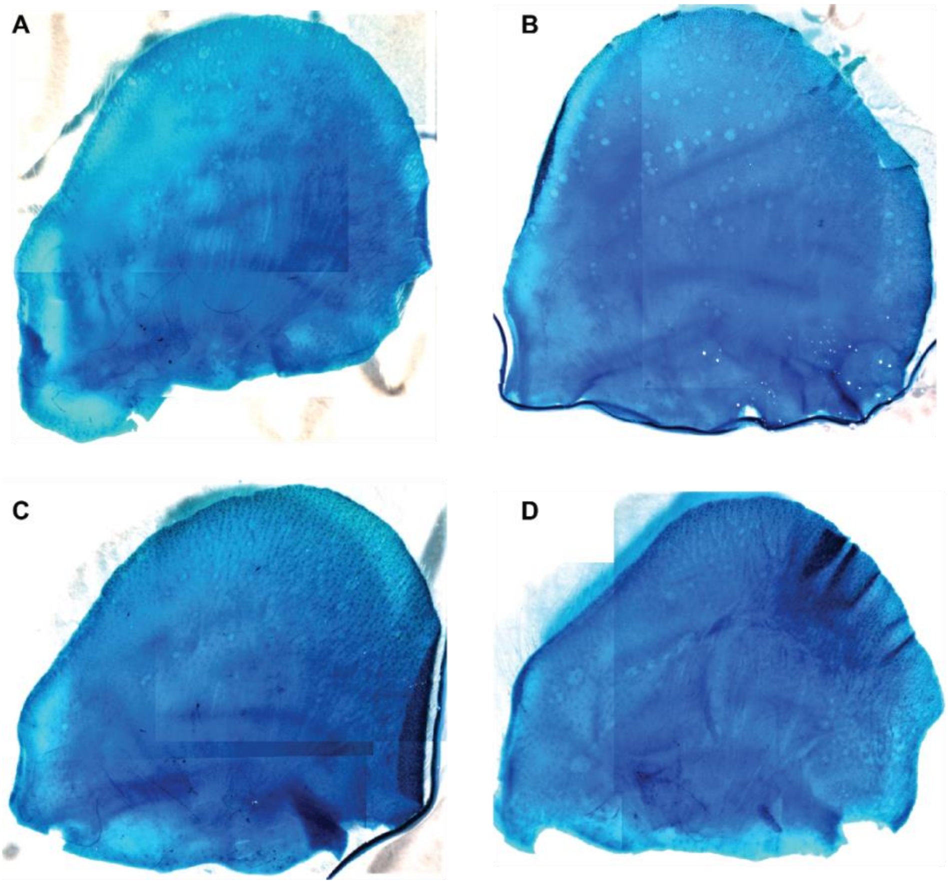


**Supplemental Figure 1:** (A-D) Partially intact adult rodent ears stained with alcian blue. Ears were collected from (A) wildtype Swiss webster *Mus*, (B) wildtype *Acomys*, (C) *Bmp5+/-* Mus, and (D) *Bmp5-/- Mus*. Dorsal Tissue was removed from all ears prior to alcian blue staining and imaging.

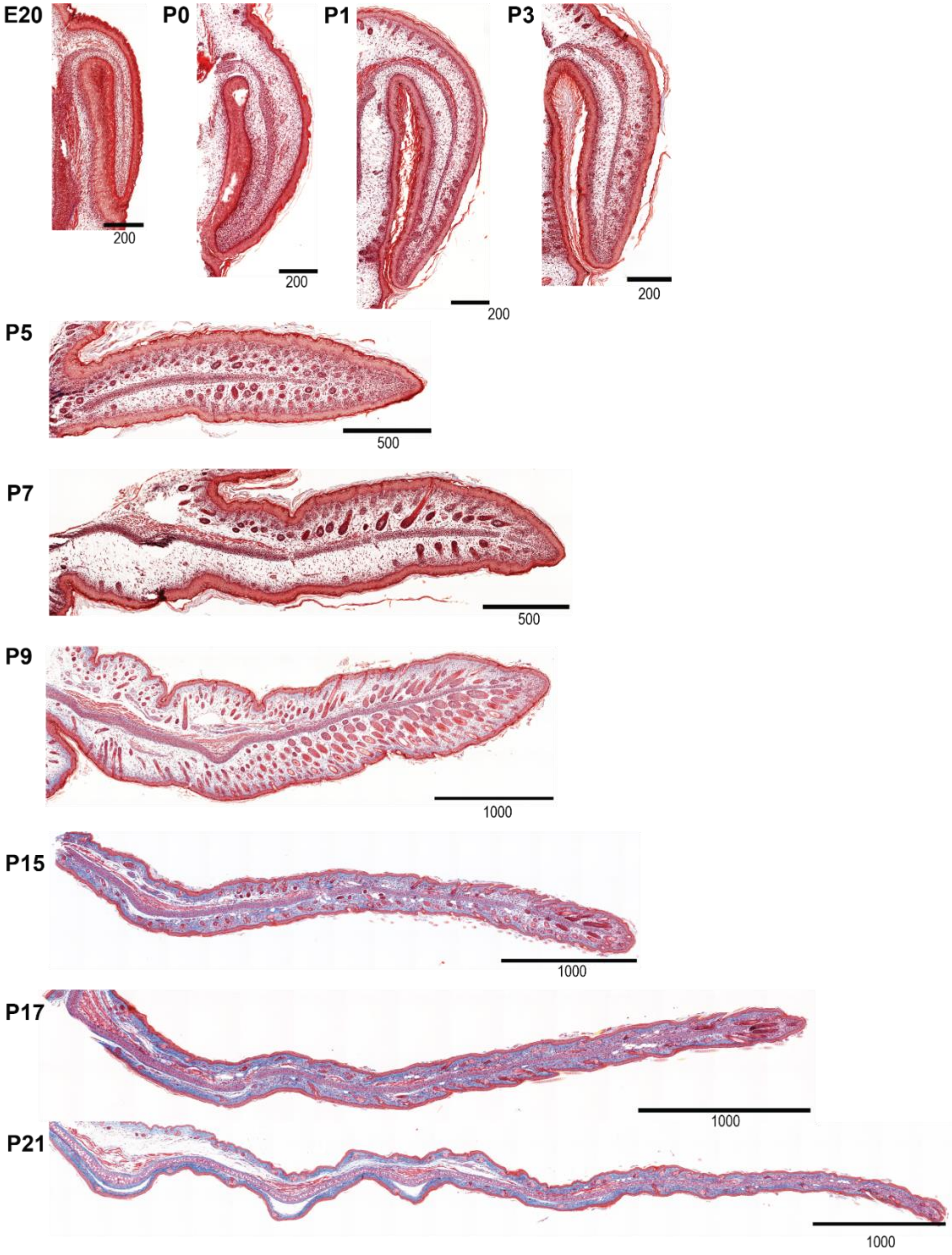


**Supplementary Figure 2:** Mus whole ear development time course. Representative pictures of wildtype Swiss webster *Mus* ear pinna at various developmental stages. Scale bar length in µm is indicated below bars.

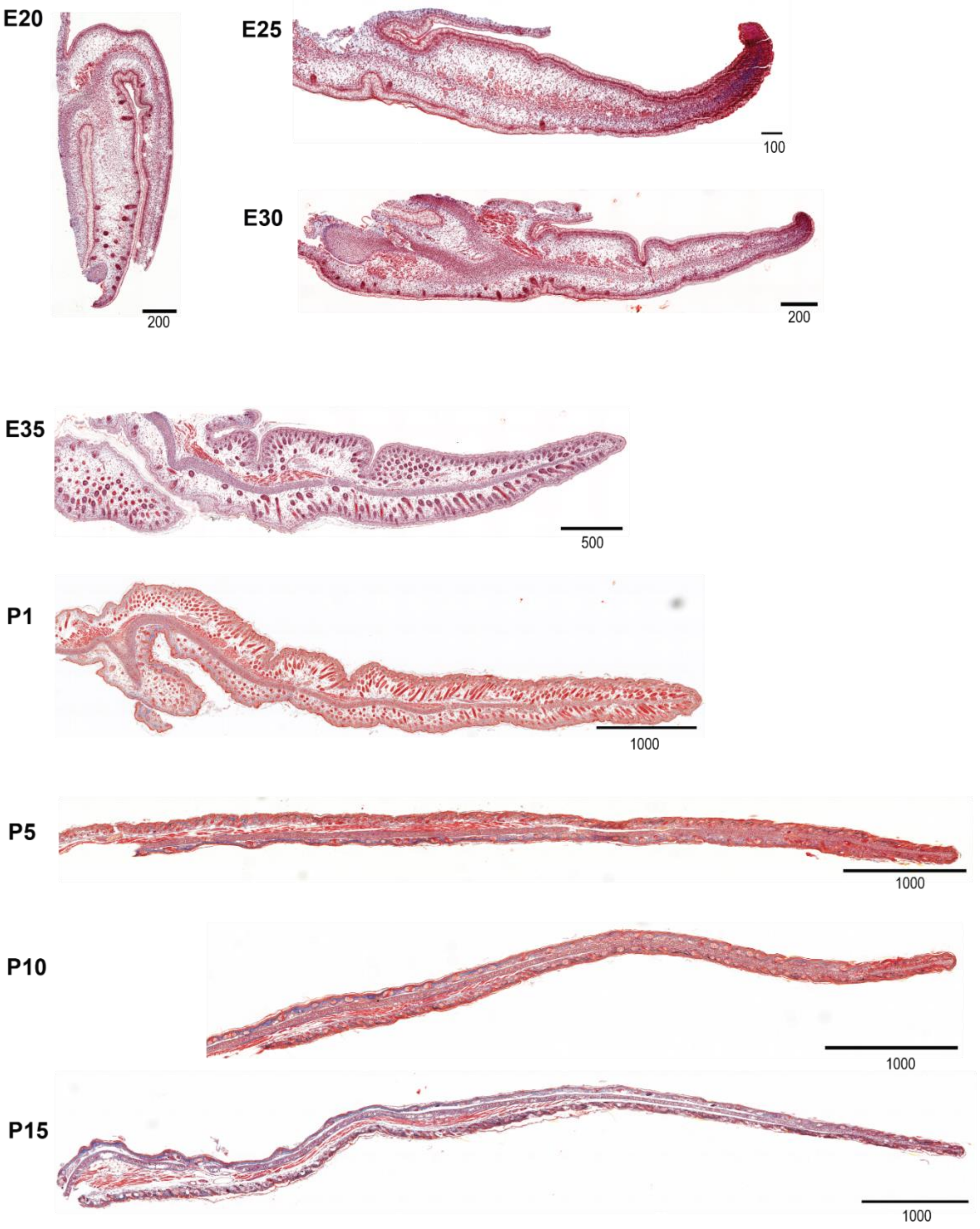


**Supplementary Figure 3:** *Acomys* whole ear development time course. Representative pictures of wildtype *Acomys* ear pinna at various developmental stages. Scale bar length in µm is indicated below bars.

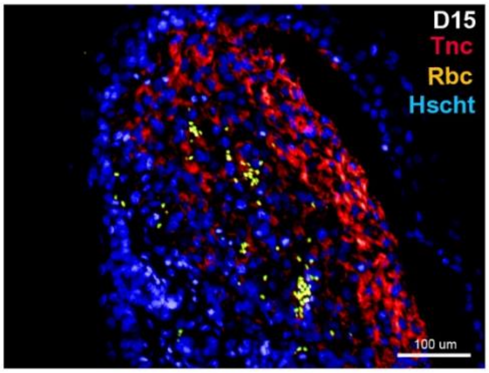


**Supplementary Figure 4:** Positive control Tenascin-C staining in *Acomys*. Tissue sample is the proximal portion of the regenerating ear pinna 15 days after 4mm biopsy excision injury through the center of the pinna.

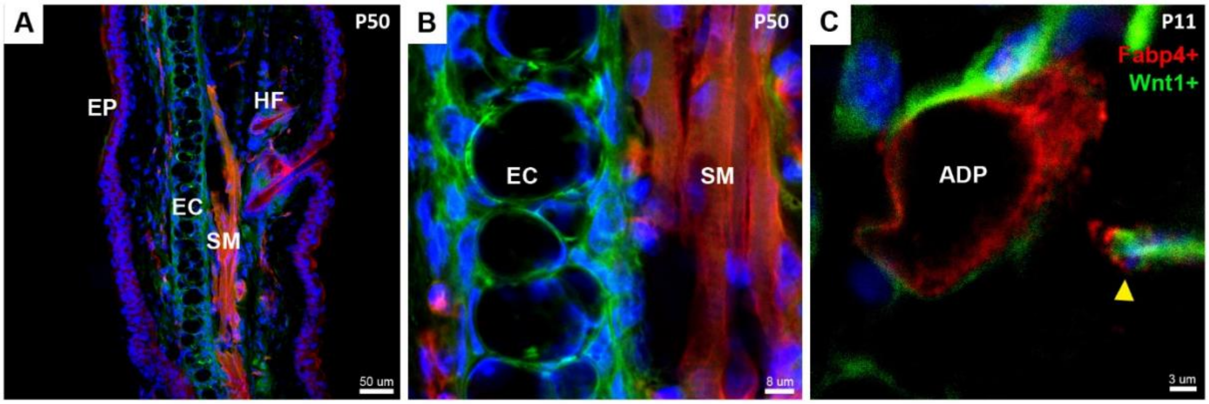


**Supplementary Figure 5:** Neural crest lineage tracing was performed by crossing male *Wnt1-Cre* driver mice to female $ROSA^{mT/mG}$ reporter *Mus*. (A) Merged *Wnt1+* (GFP), *Wnt1-* (tdTomato) and Nuclei (Hoescht) fluorescence at P50. Epidermis (EP), hair follicles (HF), elastic cartilage (EC) and skeletal muscle (SM) are shown. (B) High magnification of elastic cartilage and skeletal muscle. (C) Fabp4+ immunostained adipocytes (ADP) at P11. Yellow arrow indicates portion of an adipocyte that is out of plane.

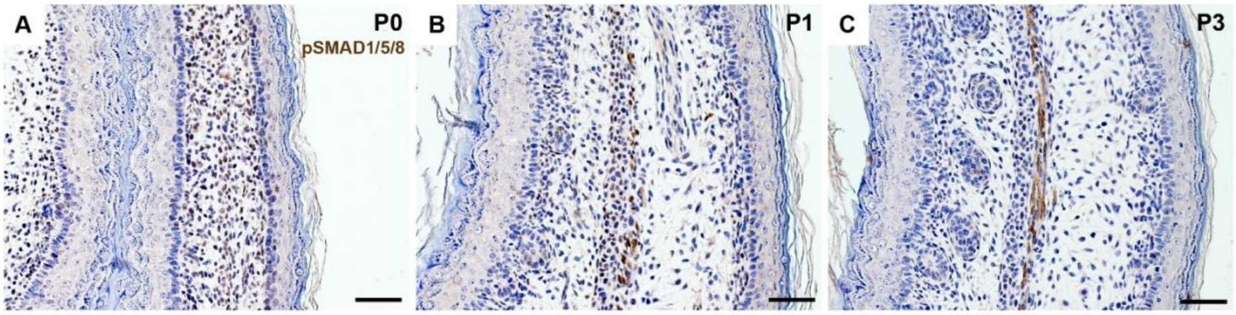


**Supplementary Figure 6:** Immunostaining for phosphosmad (pSmad)1/5/8 in wild-type *Mus* at P0 (A), P1 (B) and P3 (C). pSMAD 1/5/8 expression is marked by brown stained nuclei and observed in pre-cartilaginous condensations at P0 and P1 but absent at P3 where it is instead expressed in the developing muscle. Scale bars are 100 μm.

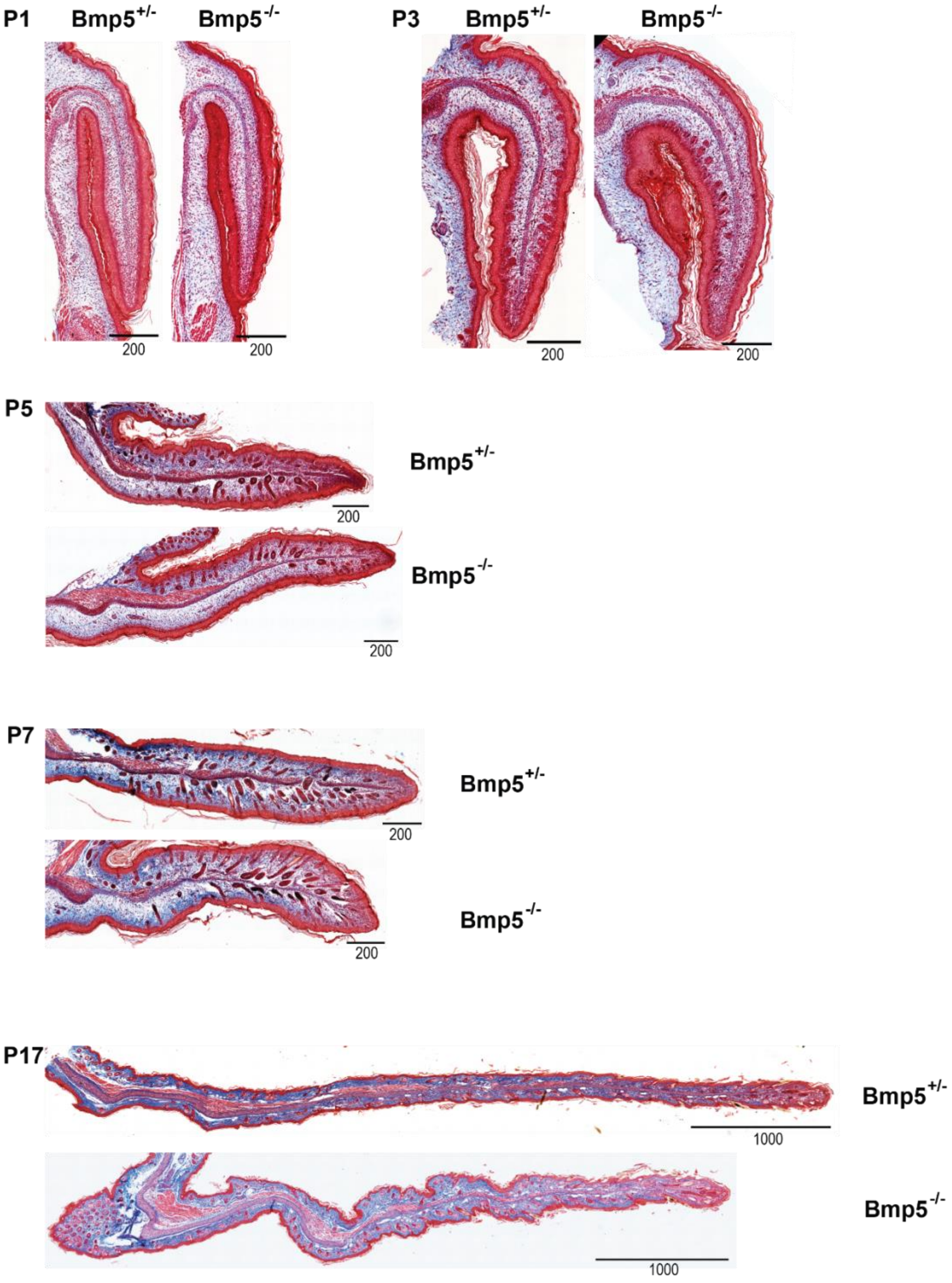


**Supplementary Figure 7:** *Bmp5*+/- and -/- *Mus* whole ear development time course. Representative pictures of Bmp5 heterozygous and Bmp5 null *Mus* ear pinna at various developmental stages. Scale bar length in µm is indicated below bars.

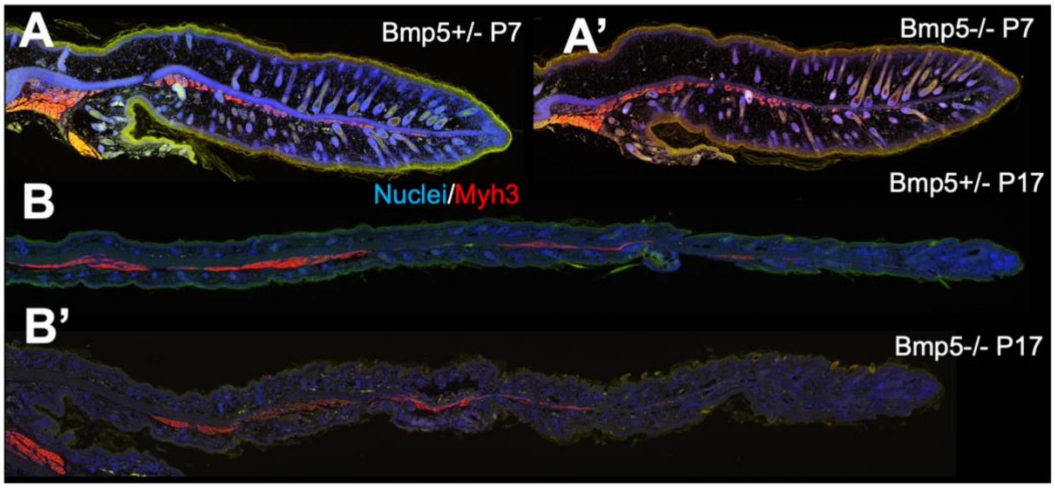


**Supplemental Figure 8:** Histological comparison of *Bmp5*+/- and -/- *Mus* ears with Myh3 immunostaining at P7 and P17.

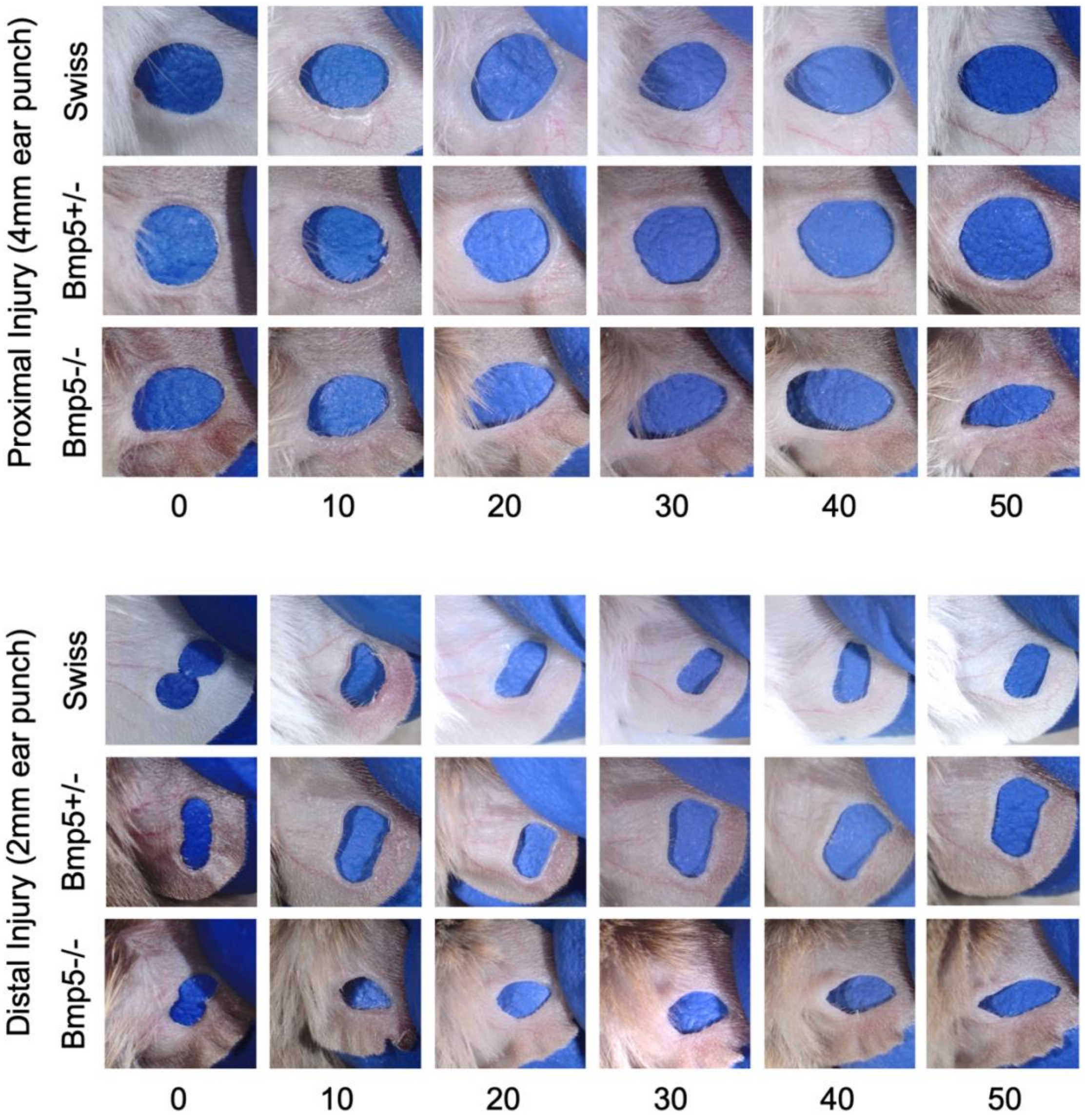


**Supplemental Figure 9:** Representative ear wounds of wildtype (Swiss webster), *Bmp5*+/-, and *Bmp5*-/- *Mus* ears after proximal 4mm or distal 2mm full thickness ear punches over 50 days of healing.
